# Overexpression of the schizophrenia risk gene C4 in PV cells drives sex-dependent behavioral deficits and circuit dysfunction

**DOI:** 10.1101/2024.01.27.575409

**Authors:** Luke A. Fournier, Rhushikesh A. Phadke, Maria Salgado, Alison Brack, Jian Carlo Nocon, Sonia Bolshakova, Jaylyn R. Grant, Nicole M. Padró Luna, Kamal Sen, Alberto Cruz-Martín

## Abstract

Fast-spiking parvalbumin (PV)-positive cells are key players in orchestrating pyramidal neuron activity, and their dysfunction is consistently observed in myriad brain diseases. To understand how immune complement dysregulation – a prevalent locus of brain disease etiology – in PV cells may drive disease pathogenesis, we have developed a transgenic mouse line that permits cell-type specific overexpression of the schizophrenia-associated complement component 4 (*C4*) gene. We found that overexpression of mouse *C4* (*mC4*) in PV cells causes sex-specific behavioral alterations and concomitant deficits in synaptic connectivity and excitability of PV cells of the prefrontal cortex. Using a computational network, we demonstrated that these microcircuit deficits led to hyperactivity and disrupted neural communication. Finally, pan-neuronal overexpression of *mC4* failed to evoke the same deficits in behavior as PV-specific *mC4* overexpression, suggesting that *C4* perturbations in fast-spiking neurons are more harmful to brain function than pan-neuronal alterations. Together, these results provide a causative link between *C4* and the vulnerability of PV cells in brain disease.

## INTRODUCTION

Cortical parvalbumin (PV)-positive fast-spiking cells are a distinct class of inhibitory neurons characterized by their expression of the Ca^2+^-binding protein, PV (1–3). Their unique biophysical properties allow them to drive potent, precise inhibition, effectively controlling the temporal dynamics of the excitatory and inhibitory inputs (4–6) that support critical brain functions (7–9). At a network level, PV cells are responsible for generating and regulating gamma oscillations (10), 30-80 Hz rhythmic fluctuations in brain activity that correlate with cognitive performance (11–13) and are impaired in anxiety disorders (14), schizophrenia (SCZ) (15,16), and Alzheimer’s Disease (AD) (17,18). Besides controlling the temporal dynamics of excitation and inhibition and orchestrating oscillatory activity, PV neuron activity tightly regulates cortical maturation during critical developmental windows (19–21).

Despite significant progress in understanding PV cell function in the healthy and diseased brain, it remains to be determined how specific genetic alterations associated with neuropsychiatric disorders lead to the dysfunction of inhibitory microcircuits. It is also unclear whether particular circuitry or synaptic inputs underlying the function of PV cells are susceptible to disease-associated genetic alterations. For example, in post-mortem tissue of patients with SCZ, the density of excitatory synapses is decreased on PV cells with concomitant downregulation of PV and other inhibitory markers (22–25), suggesting that excitatory drive to fast-spiking cells is compromised in this brain disorder. Moreover, SCZ-associated genetic alterations have been found to disrupt the molecular machinery underlying feed-forward excitatory inputs to PV neurons (26–28), suggesting these particular connections are susceptible to genetic perturbations.

The association between the Major Histocompatibility Complex (MHC) and SCZ could provide a link between immune dysfunction and the disruption of molecular mechanisms that regulate the wiring of synaptic circuits (29–32). In support of this, Sekar et al. (33) showed that mice that lack *C4b* (*mouse C4*, *mC4*), which in humans is harbored in the MHC locus, exhibit deficits in the developmental refinement of retinogeniculate synapses. Furthermore, Comer et al. (34) demonstrated that increasing levels of the human (*C4A*) and mouse *C4* homologs in layer (L) 2/3 pyramidal neurons (PYRs) of the medial prefrontal cortex (mPFC) – a brain region associated with the pathology of SCZ (35–37) and other neuropsychiatric conditions (38–40) – led to pathological synaptic loss during early postnatal development and social behavioral deficits in mice. This research suggests a link between immune dysfunction and brain disorders, particularly SCZ, through the role of the MHC and its impact on synaptic development and plasticity.

As a consequence of their unique properties and role in controlling network function, PV cells exhibit high metabolic demands, which make them vulnerable to oxidative stress and neuroimmune dysregulation (41–45); these pathological processes are linked to brain disorders (46–48). Therefore, determining the molecular pathways through which immune imbalances can impact PV neuron function offers significant potential for unraveling neurodevelopmental disease etiology. Despite this, there is a notable gap in the availability of models to determine how neuroimmune dysfunction alters specific brain circuitry. In the same vein, alterations in the complement pathway have been linked to the pathology of brain disorders (49). However, it remains an open question whether particular brain cell types are especially vulnerable to complement dysfunction.

Here, we developed and validated a novel mouse line that permits cell-type specific overexpression of *mC4* (mC4-OE). Utilizing this unique knock in (KI) transgenic mouse, we demonstrate that increased *mC4* levels in PV cells (PV-mC4-OE) drive pathological anxiety-like behaviors in male, but not female mice. In both sexes, PV-mC4-OE led to changes in a subclass of social behavior, indicating that elevated expression of this immune gene in fast-spiking cells disrupts the circuitry governing social behaviors. We used electrophysiology to show that excitatory and inhibitory inputs to mPFC PV cells are altered with increased levels of *mC4* in male, but not female mice, mirroring the sexually dimorphic anxiety-like behavior.

Elevated *mC4* in PV neurons had differing impacts on cortical cell excitability: hypoexcitability in male fast-spiking cells and PYRs, as opposed to enhanced excitability in female fast-spiking cells. In contrast to PV-cell-specific effects of mC4-OE, increased *mC4* levels in all neurons had no effects on anxiety-like behavior. This implies that *C4* perturbations in PV cells are more harmful to brain function than pan-neuronal alterations. Finally, using a computational model, we demonstrated that deficits in the PV-mC4-OE-driven inhibitory microcircuit disrupt model pyramidal neuron communication and cause hyperexcitability within the male model network, illustrating how the interaction of cellular and synaptic traits generates complex sex-dependent neural network deficits in disease states.

In summary, our results show that *mC4* dysregulation in PV neurons led to alterations in anxiety-like behavior in male mice, which were associated with mPFC circuit dysfunction. Our results provide crucial insights into the molecular interplay between cell-type-specific increased levels of *mC4*, defects in mPFC circuitry, and abnormal behavior associated with the prefrontal cortex.

## METHODS

### Ethics Statement and Animals

All experimental protocols were conducted according to the National Institutes of Health (NIH) guidelines for animal research and were approved by the Boston University Institutional Animal Care and Use Committee (IACUC; protocol #2018–539). All mice were group-housed (2-4 animals/cage). The light/dark cycle was adjusted depending on the behavioral task (see below). Unless otherwise stated, food and water were provided *ad libitum* to all mice. Experimental offspring were reared in the cage with the dam until weaning at postnatal day (P) 21. Stimulus CD-1 (Charles River Laboratories, strain code: 022, RRID:IMSR_CRL:022) mice 3-5 weeks of age were used in social assays. During experiments and analysis, the experimenter was blinded to mouse genotype and experimental conditions whenever possible.

#### Generation of the mC4-KI mouse

Generation of the mC4-KI mouse was accomplished using Cyagen/Taconic services (Santa Clara, CA). Briefly, the “adenovirus SA-Frt-CAG promoter-Frt-loxP-3*polyA-loxP-Kozak-mouse C4b CDS-polyA” cassette was cloned into intron 1 of ROSA26. The homology arms were generated by PCR using BAC clones as templates. C57BL/6 ES cells were used for gene targeting. In the targeting vector, the positive selection marker (Neo) is flanked by SDA (self-deletion anchor) sites. Diphtheria toxin A (DTA) was used for negative selection. Targeted ES cells were injected into C57BL/6J albino embryos, which were then re-implanted into CD-1 pseudo-pregnant females. Founder animals were identified by their coat color, and their germline transmission was confirmed by breeding with C57BL/6J females and subsequent genotyping of the offspring. Male and female heterozygous targeted mice were generated from clone 1F3 and were bred to establish a colony. Lastly, the constitutive KI cell allele was obtained after Cre-mediated recombination (see *Breeding* section).

#### Breeding

C57BL/6J (Jackson Laboratory, strain #: 000664, RRID:IMSR_JAX:000664) mice were paired with heterozygous mC4-KI transgenics of the same genetic background.

Genotyping (Transnetyx) was used to plan breeding schemes and identify specific genotypes. Heterozygous mC4-KI transgenics were paired with homozygous PV-Cre mice (Jackson Laboratory, B6.129P2-Pvalb^tm1(cre)Arbr^/J, strain #: 017320, RRID:IMSR_JAX:017320). Our breeding scheme generated mice that inherited the floxed mC4-KI allele, and thus overexpress (OE) *mC4* in PV cells (PV-mC4-KI, or simply KI), and wild type littermates, which were used as control mice (PV-mC4-WT, or simply WT).

To OE *mC4* in all neurons, heterozygous mC4-KI transgenic mice were paired with homozygous BAF53b Pan-neuronal-Cre (Jackson Laboratory, STOCK Tg(Actl6b-Cre)4092Jiwu/J, Strain #:027826, RRID:IMSR_JAX:027826, (50)) mice. Breeding these two mouse lines yielded mice that inherited the floxed mC4-KI allele, and thus OE *mC4* in all neurons (PanN-mC4-KI), and wild type control littermates (PanN-mC4-WT).

Our breeding scheme ensured that all experimental mice carried the cre recombinase gene to control for its effects. PV-mC4-KI or PanN-mC4-KI mice did not exhibit any gross brain abnormalities, indistinguishable from controls. Additionally, they had similar weights to their controls, suggesting that besides the described cellular and behavioral deficits, these mice were otherwise healthy and had no significant defects.

A strength of the mC4-KI transgenic mouse is that more moderate levels of mC4-OE can be driven in a cell-specific manner by crossing the mC4-KI mice to Flp transgenic mice, which substitutes the strong CAG promoter for the weaker ROSA26 promoter (51,52), allowing the effects of more moderate transgene expression to be studied.

### Multiplex fluorescence *in situ* hybridization

#### Tissue preparation and staining

For multiplex fluorescence *in situ* hybridization (M-FISH) experiments, brains of PV-mC4-WT and PV-mC4-KI mice P21-22 or P58-65 were extracted and immediately embedded in O.C.T. (Tissue-Tek, 4583), flash-frozen on dry ice, and stored in - 80°C until being cut. Prior to slicing, brains were moved to -20°C for 30 min. Slices were cut on a Leica CM 1800 cryostat at 16 μm at -16 to -19°C and adhered directly onto microscope slides (Fisher Brand Superfrost Plus, #1255015), which were then stored in -80°C until ready for M-FISH (<1 week). M-FISH experiments were then performed as directed by the commercially available kit (RNAScope, Advanced Cell Diagnostics), from which all probes and reagents were purchased. Fluorescent probes for *mC4* (Mm-C4b, 445161), parvalbumin (Mm-Pvalb-C2, 421931-C2), and somatostatin (Mm-Sst-C3, 404631-C3) were used. To confirm cell bodies in the slice, nuclei were stained with RNAScope DAPI (320858). After staining, the slices were mounted with ProLong Diamond Antifade Mountant (ThermoFisher, P36961). Each round of M-FISH performed contained tissue from both PV-mC4-WT and PV-mC4-KI mice.

#### Imaging and analysis

M-FISH images were acquired at 40x on a confocal laser scanning microscope (Nikon Instruments, Nikon Eclipse Ti with C2Si^+^ confocal), controlled by NisElements (Nikon Instruments, 4.51) including four laser lines (405, 488, 561, and 640 nm), at a step size of 0.4 μm for nearly the entire thickness of the tissue slice (16 μm). For each round of M-FISH, consistent imaging parameters were used. Tissue slices imaged and analyzed belonged to mPFC divisions (prelimbic (PrL), infralimbic (IL), and anterior cingulate (AC) cortices) of the mouse brain. Images predominantly included L2/3, though deeper cortical layers were also included in the region of interest (ROI). For analysis, a maximum intensity projection of each z-stack was made (ImageJ, National Institute of Health, Bethesda, Maryland) and transferred into CellProfiler (53,54) (Broad Institute). Cells were identified and segmented using DAPI, and the contour was expanded by 10 pixels (approximately 3.1 µm) to capture the majority of *Pv*, *Sst*, and *mC4* puncta surrounding the nucleus. Cells were classified as *PV* or *SST* cells if their expanded contour contained an empirically-derived minimum of 13 or 10 *Pv*-or *Sst*-positive puncta, respectively. Once identified as a *PV*, *SST*, or non-*PV*/non-*SST* DAPI+ ‘Other’ cell, CellProfiler was used to quantify the number of *mC4-*positive puncta within each contour (i.e., within each cell).

### PV cell density

#### Perfusion and immunohistochemistry

Mice P55-74 were anesthetized with a 4% isoflurane-oxygen mixture (v/v) and perfused transcardially with phosphate-buffered saline (PBS, Gibco, Life Science Technologies, 70011044) followed by 4% paraformaldehyde (PFA) in PBS. Extracted brains were further fixed in PFA for 24 h before being transferred to a 30% (w/v) sucrose solution and stored at 4°C. Brain slices were cut at 40 μm on a freezing stage sliding microtome (Leica SM2000) and stored in PBS. From each mouse, two brain slices were selected for immunostaining: one at approximately Bregma +1.98 mm and another at Bregma +0.98 mm along the anterior-posterior (A-P) axis. Slices were first blocked and made permeable in a solution containing 10% donkey serum (Sigma-Aldrich, S30-100ML) and 1% Tritonx100 (Sigma-Aldrich, X100-100ML) in PBS. Next, after applying the primary antibody (rabbit anti-PV, Abcam, ab11427) at a 1:250 dilution, slices were placed on a shaker at 4°C for 48 h. Slices were then washed 4 x 15 min with 0.025% Tritonx100 in PBS. Next, the secondary antibody was applied (anti-rabbit STAR RED, Abberior, STRED) at a 1:500 dilution and returned to the shaker for 48 h at 4°C. Slices were then washed 4 x 15 min in PBS, and mounted onto 1 mm microscope slides (Globe Scientific, #1324) with DAPI Fluoromount (Thermo Fisher Scientific, Cat. #: 00-4959-52).

#### Imaging and Analysis

Cell density imaging was acquired (laser lines 405 and 640 nm) at a step size of 1 μm for nearly the entire thickness of the tissue slice (40 μm). For each animal, the mPFC was imaged at 20x in both the slice at Bregma +1.98 mm (6 ROIs total, 3/hemisphere, that include the PrL, IL, and AC cortices) and the slice more posterior at Bregma +0.98 mm (4 ROIs, 2/hemisphere, all AC). Consistent imaging parameters were maintained across all imaging sessions. Images were analyzed as TIFFs in ImageJ and compared to a brain atlas to identify brain regions (55,56). The intensity value in the PV channel for each ROI (in a brain slice) and the average background signal for each brain slice were quantified. To binarize cells as PV-positive, we calculated an intensity threshold and classified cells as PV-positive if their intensity value was higher than this threshold. These data were used to calculate the number of PV cells that were positive. To calculate density, we determined the area (excluding L1) of the ROI in which PV cells were counted and calculated the 3D volume (in mm^3^) by multiplying the 2D area of each slice by the depth of the tissue imaged (the Z-stack).

### Behavior

#### General experimental conditions

P40-60 mice were used for all behavioral assays, group housed in sets of 2-3 mice per cage. Mice were used in either (1) a series of anxiety-related assays or (2) a series of sociability assays. The specific sequence of anxiety-related assays that mice were exposed to was consistent across all mice and proceeded in the following order: open field (OF) and elevated zero maze (EZM, performed on the same day), light-dark box (LDB), and novelty-suppressed feeding (NSF) (Novel arena and cage NSF, performed back-to-back).

Similarly, the specific sequence of sociability assays that mice were exposed to was also consistent across all mice: Object and juvenile interaction (performed immediately back-to-back for each mouse, see *Object and social interaction*) was followed by the three-chamber sociability assay.

Seven days prior to the first day of handling (see *Handling* below), mice were genotyped (Transnetyx), transferred to a fresh cage, and placed with a *Do Not Handle* card to minimize human handling and stress. Mice were reliably identified using a set of ear hole-punches throughout behavioral experiments.

All behavioral assays were performed at a similar time of the day. Mice used in anxiety-related assays were reared on a 12 h light/dark cycle with lights on at 7 AM and lights off at 7 PM, and assays were performed under white light (Adorama, 13” Dimmable LED Ring Light). The intensity of light used for each assay was consistent each time the assay was performed, but varied dependent on the assay (see specific assays below). Each day, the lux was measured and adjusted to the appropriate level for the assay being performed (Dr.meter LX1330B). Mice used in sociability assays were reared on an inverted light/dark schedule, with lights on at 7 PM and lights off at 7 AM, and assays were performed under red light (Amazon Basics 60W Equivalent, Red) to minimize the stress-inducing effects of bright white light and to remain consistent with their inverted light/dark schedule. Behavioral assays in a given series were always separated by at least two days but never more than four days. On all days of behavioral experiments and handling, mice were retrieved from the facility and left in the behavior room to acclimate to the environment for at least 1 h. Once all mice in a cage completed the assay, all were returned to their original home cage.

Acquisition of behavior data for all anxiety-related assays and the three-chamber assay was recorded using Logitech C922x Pro Stream Webcams at 30 frames per second (fps) via the open-source UCLA miniscope software (57). Acquisition of object and juvenile interaction data was recorded at 30 fps using a Teledyne Flea3 USB3 camera (Model: FL3-U3-13E4C-C: 1.3 MP, e2v EV76C560) via an in-house, python-based, open-source video acquisition software, REVEALS (https://github.com/CruzMartinLab/REVEALS) (58).

Throughout all components of behavioral assays and handling, gloves and a lab coat were worn. Gloves, behavioral arenas, and any relevant objects or cups used during the assay were sprayed with 70% ethanol in between handling mice or between each new mouse performing a given assay.

All behavioral assays were performed blind to condition, and analysis was performed via custom code written in MATLAB (MathWorks) (see *Behavior Analysis* below).

#### Handling

The first assay of each series was always preceded by three consecutive days of handling. Mice involved in anxiety-related assays were handled under standard, ambient room lighting (270 lux), and mice involved in sociability assays were handled under red light.

### Anxiety-related Assays

Anxiety-inducing arenas were custom-made as described in Johnson et al. (59).

#### Open field (OF)

Mice were placed in the center of a custom-made OF, a (45 × 45 × 38 cm, length x width x height) black arena under 200 lux of white light, and were free to explore for 10 min. The OF was used to measure locomotion by measuring the distance traveled by the mice.

#### Elevated zero maze (EZM)

The EZM is an elevated (63 cm) circular platform with a 5 cm track width and diameter of 60 cm. It is comprised of two closed arms with a wall height of 14 cm and two open arms that lack walls. The EZM was run under 200 lux of white light for 10 min.

#### Light-dark box (LDB)

The LDB uses the frame of the OF, but inserted into the OF is a black divider (45 x 38 cm, length x height) that divides the OF into two distinct zones 1/3 and 2/3 the width of the OF, but features a small passage-way at the bottom (7.6 x 7.6 cm, width x height) to allow the mice to move freely between zones. Over the smaller zone (45 x 15 cm, length x width) is a black lid that blocks all light: this is the dark zone. Because there is no lid over the remaining 2/3 of the OF (45 x 30 cm, length x width), this is the light zone. The LDB was run under 300 lux of white light for 10 min.

#### Novelty-suppressed feeding (NSF)

24 h before the start of experimentation, mice were transferred to a clean cage that possessed no food. During the assay, mice were placed in a novel, open arena (50 x 35.5 x 15 cm, length x width x height) with a single, fresh food pellet strapped down in the center of the arena with a rubber band. The experimenter watched the live video feed to observe when the mouse traveled to the center of the maze and began feeding on the food pellet, which concluded the assay. Simple investigation or sniffing of the pellet was not considered feeding. The NSF was the only hand-scored assay, and was done so by a trained, blinded experimenter. To measure the latency to feed, the researcher watched the trial back and determined the exact frame that the mouse first bit the pellet. In the rare event that the mouse did not eat the pellet in the time allotted (10 min recording), that mouse was excluded from the NSF analysis. The NSF was run under 200 lux of white light.

#### Cage NSF

Immediately following the NSF, mice were placed in a fresh cage with approximately 10 food pellets placed in one corner of the cage. Again, the latency to feed was recorded by the experimenter. The Fresh Cage NSF served to determine and verify the anxiety-inducing nature of the arena relative to the more familiar environment of a standard mouse cage.

#### Z-Anxiety quantification

Z-Anxiety quantification was adapted from Guilloux et al. (60). For each individual assay, the mean (*µ*) and standard deviation (*σ*) of the relevant metric for that assay (e.g., EZM: time spent in open arms) for all WT animals was calculated. For any given mouse in any single assay, whose performance in that assay (e.g., EZM: time spent in open arms) is given by *x*, the z-score is the following:

To be consistent with a positive z-score being indicative of increased anxiety, the sign of the z-score in the EZM and LDB was multiplied by -1. In this way, less time spent in the open arms (EZM) or in the light zone (LDB), indicators of increased anxiety-like behavior, yielded positive z-score values. For the NSF, because an increased latency to feed was indicative of anxiety-like behavior, these z-scores were not multiplied by -1. Each animal’s Z-Anxiety is simply an average across all three assays, given by the following:

For the sex-separated quantification of Z-Anxiety, the only modification made was that the z-score for each assay for all males was found in reference to the WT male average and standard deviation; likewise, z-scores for females were made in reference to the WT female average and standard deviation for each assay.

### Sociability Assays

#### Object and social interaction

Mice were habituated to the empty, clear arena (46 x 23 cm). Once 120 s elapsed, a single novel object made from two glued 6-well plates was temporarily secured at one end of the arena with a magnet. Mice were free to explore this object for another 120 s. After this time elapsed, the novel object was removed, and was immediately replaced by a novel, juvenile (3-5 weeks old), sex-matched CD-1 stimulus mouse. These mice were then free to interact with one another unencumbered for 120 s.

#### Three-chamber sociability assay

One day prior to the three-chamber assay, mice were habituated to the three-chamber apparatus (three 46 x 23 cm chambers connected by passageways 10 cm wide). These mice were free to explore the entire three-chamber arena with an empty wire-mesh pencil cup in each of the two end chambers for 5 min. On that same day, stimulus CD-1 mice were habituated to being placed underneath these wire-mesh pencil cups for 5 min. The next day, the three-chamber assay was performed as follows. A novel, juvenile, sex-matched CD-1 stimulus mouse was placed underneath a cup in the chamber at one end of the arena, and an empty cup was placed in the chamber at the other end of the arena. Weights sat on top of the cups to ensure that the CD-1 stimulus mouse was secure in the cup and that the experimental mouse would not move either cup. To begin the assay, dividers were placed in the three-chamber passageways to block movement between chambers, and the experimental mouse was placed in the center chamber. After the mouse explored the center chamber for 120 s, the inserts were removed, and the mouse was free to explore the entire three-chamber arena, as well as the empty cup and mouse cup for 10 min. The chambers in which the mouse cup and empty cup were placed was alternated randomly across mice. The percent time spent in each of the three chambers was scored.

### Behavior Analysis

In all behavioral assays (except for the NSF and Fresh Cage NSF), mice and relevant behavioral arena components were tracked using DeepLabCut (DLC) (61). To ensure the accuracy of tracking by DLC, a random sampling of videos from each day of experimentation were inspected. Next, a trained experimenter watched annotated videos to verify consistent tracking of fiducial points. Fiducial points included the snout, the nose bridge, the head, left and right ears, and five points that ran along the sagittal axis of the mouse body from the neck to the base of the tail. All corners of all arenas were labeled, as well as any relevant features of the arenas, including corners of objects and cups, and the thresholds separating the open and closed arms of the EZM. To calculate the interaction times, binary behavior matrices (vectorized behavior) indicating the location of the relevant key points of the mouse with respect to relevant key points of the arena (e.g., head of mouse and corner of object) were created using custom MATLAB scripts, available at (https://github.com/CruzMartinLab/PV-mC4_Project/tree/main/behavior_code).

### Neonatal viral injections

To genetically tag and identify PV cells in electrophysiology experiments, P1-3 pups were injected with 360 nL of AAV1-FLEX-tdTomato (titer: 2.5×10^13^ vg/mL, Addgene #28306) per cortical hemisphere. Borosilicate pipettes (BF150-117-10, Sutter Instrument Co., Novato, California) were pulled to a fine tip (approximately 3-15 μm) and back-filled with mineral oil and inserted into the Nanoject Injector (Drummond, Nanoject II, 3-000-204/205A/206A). After cutting the tip of the pipette and emptying roughly half of the mineral oil, the pipette was filled with virus solution from the open tip. Prior to injection, pups were anesthetized via a cold-plate (approximately 15 min) and remained on the cold surface of an ice pack during injection to ensure continued anesthesia throughout the entire process. The mPFC was targeted along the anterior-posterior axis and hit consistently using empirically derived landmarks and the Allen Brain Atlas (55). From here, the tip of the pipette was moved medially into position immediately adjacent to the midline. Injections at several depths were made to ensure effective labeling of the entire dorsal-ventral depth of mPFC target sub-regions. Fine spatial navigation of the tip was made using a stereotax (Kopf Instruments, Tujunga, California). Post-injection, pups recovered in a plastic chamber that was placed on top of a heated blanket. Pups were returned to the dam once fully recovered.

### Electrophysiology

#### Acute slice preparation and recording

Mice (P40-62) were anesthetized with a 4% isoflurane-oxygen mixture (v/v) and perfused intracardially with ice-cold Perfusion/Slicing artificial cerebrospinal fluid (P/S-aCSF) bubbled with 95% O_2_/ 5% CO_2_ containing the following (in mM): 3 KCl, 26 NaHCO_3_, 1.25 NaH_2_PO_4_, 212 sucrose, 10 glucose, 0.2 CaCl_2_, and 7 MgCl_2_ (300-310 mOsm). Thirty min before slicing, 200 mL of P/S-aCSF was transferred to -20°C until turned to a slushy consistency. Coronal slices 300-μm thick were cut in this slushy P/S-aCSF using a VT1000 S (Leica) vibratome and were then transferred to a Recording aCSF (R-aCSF) solution bubbled with 95% O_2_/ 5% CO_2_ containing the following (in mM): 125 NaCl, 2.5 KCl, 25 NaHCO_3_, 1.4 NaH_2_PO_4_, 16 glucose, 0.4 Na-ascorbate, 2 Na-pyruvate, 2 CaCl_2_, and 1 MgCl_2_ (300-310 mOsm). Slices were incubated in this R-aCSF for 30 min at 35°C before being allowed to recover at room temperature for 1 h prior to recording.

Whole-cell voltage- and current-clamp recordings were performed in Layer (L) 2/3 of the PrL, IL, and AC cortex divisions of the mPFC (34). For all recordings, tdTomato-positive PV cells were identified using a Prior Lumen 200 Light Source (Prior Scientific) and a CMOS camera (Rolera-Bolt-M-12; 1.3 MP, Mono, 12-BIT, Uncooled, QImagingBolt) mounted on an Olympus BX51WI microscope (Olympus America, Inc.). Pyramidal neurons (PYRs) were identified based on morphological and electrophysiological properties. All recordings were performed at 29-31°C. Signals were recorded with a 5X gain, low-pass filtered at 6 kHz, and digitized at 10 kHz using a patch-clamp amplifier (Multiclamp 700B, Molecular Devices). Nearly all recordings were made using 3-5 MΩ borosilicate pipettes (Sutter, BF-150-117-10). Series (R_s_) and input resistance (R_in_) were monitored throughout the experiment by measuring the capacitive transient and steady-state deflection in response to a −5 mV test pulse, respectively. Liquid junction potentials were calculated and left uncompensated.

#### Miniature post synaptic currents (mPSCs)

For recording miniature excitatory post synaptic currents (mEPSCs), borosilicate pipettes were filled with an internal recording solution that contained the following (in mM): 120 Cs-methane sulfonate, 8 NaCl, 10 HEPES, 10 CsCl, 10 Na_2_-phosphocreatine, 3 QX-314-Cl, 2 Mg^2+^-ATP, and 0.2 EGTA (292mOsm, adjusted to pH 7.3 with CsOH). PV cells and PYRs were voltage clamped at −70 mV in the presence of tetrodotoxin (TTX, 1 μM, Tocris) and picrotoxin (PTX, 100 μM, HelloBio). 6-9 mice per sex per condition from 3-4 litters were used to collect all mEPSC data for PV cells and PYRs.

For recording miniature inhibitory post synaptic currents (mIPSCs), borosilicate pipettes were filled with a high-chloride internal recording solution that contained the following (in mM): 60 Cs-methane sulfonate, 8 NaCl, 70 CsCl, 10 HEPES, 10 Na_2_-phosphocreatine, 0.2 EGTA, and 2 Mg^2+^-ATP (290 mOsm, adjusted to pH 7.3 with CsOH). PV cells and PYRs were voltage clamped at -70 mV in the presence of TTX (1 μM), CNQX (20 μM), and DL-APV (50 μM).

Because this high-chloride internal solution has a chloride reversal potential of -13 mV, mIPSCs were inward. 4-6 mice per sex per condition from 3-5 litters were used to collect all mIPSC data for PV cells and PYRs.

#### mPSC analysis

mPSCs were identified and their amplitude, frequency, rise, and decay determined using custom scripts written in MATLAB. At least 120 s were analyzed for each cell. All mPSC raw traces were first lowpass filtered in Clampfit (Molecular Devices) using a boxcar filter. Next, local minima in the trace were recognized by identifying potential synaptic events using the native *islocalmin()* MATLAB function. After these events were filtered, a series of steps were taken to remove false-positives while simultaneously limiting the number of false negatives. More specifically, we calculated a threshold based on the standard deviation of the noise of the raw trace within a 1 s temporal window to differentiate the background noise from mPSCs, thus setting an amplitude threshold. Next, a series of thresholds based on the rise and decay times were used to filter subsequent postsynaptic events. For all remaining mPSCs, amplitude is given as the difference between the baseline current value (determined using a highly smoothed line of the raw data that effectively serves as a moving baseline of the trace) at the time when the peak reaches a minimum current value and the average of the 10 points around the absolute minimum of that mPSC peak.

Frequency (in Hz) of postsynaptic events is given by the number of mPSCs per sec. Rise_10-90_ is defined as the time (ms) it takes for the mPSC to progress from 10 to 90% of the peak of that mPSC. To find the Decay_tau_, for each event, the trace from the peak of the mPSC to its return to baseline is isolated and fit to a single-term exponential. Between groups, R_s_ values for each cell type were not statistically significant.

R_s_ for PV cells (mEPSCs) were as follows (in MΩ): WT males: 16.85 ± 1.17; WT females: 17.35 ± 1.26; KI males: 16.27 ± 1.33; KI females: 15.56 ± 1.21. R_s_ for PV cells (mIPSCs) were as follows (in MΩ): WT males: 15.79 ± 1.06; WT females: 13.93 ± 1.09; KI males: 15.40 ± 1.46; KI females: 14.07 ± 0.79. R_s_ for PYRs (mEPSCs) were as follows (in MΩ): WT males: 15.94 ± 0.92; WT females: 14.81 ± 0.75; KI males: 15.07 ± 0.73; KI females: 15.00 ± 0.80. R_s_ for PYRs (mIPSCs) were as follows (in MΩ): WT males: 14.43 ± 1.05; WT females: 13.81 ± 0.92; KI males: 16.13 ± 1.34; KI females: 14.88 ± 1.26.

#### Active and passive properties

To determine the active and passive properties of PV cells and PYRs, borosilicate pipettes were filled with an internal solution that contained the following (in mM): 119 K-gluconate, 6 KCl, 10 HEPES, 0.5 EGTA, 10 Na_2_-phosphocreatine, 4 Mg^2+^-ATP, and 0.4 Na-GTP (292 mOsm, adjusted to pH 7.3 with KOH). Cells were held at -65 mV during recording, and the bath was perfused with CNQX (20 μM), DL-APV (50 μM), and PTX (100 μM).

Excitability was assessed by measuring membrane voltage changes (i.e., current-evoked Action Potentials (APs)) to a spiking protocol that applied 500 ms square current pulses to the patched cell, beginning at -250 pA and increasing in 30 pA steps to a max current injection of 470 pA. Passive properties of the patched cells were determined via a 500 ms, -20 pA square pulse that preceded the square pulse of increasing current amplitude. This protocol was run and recorded 2-3 times per cell, and final values were averaged across recordings for each cell. 5-7 mice per sex per condition from 4-5 litters were used to collect all active and passive properties data for PV cells and PYRs.

#### Active properties analysis

To quantify spike frequency (Hz), the number of spikes (temporally defined as when the rising phase of the spike crossed 0 mV) was divided by the length of the current pulse (0.5 s). Rheobase was defined as the minimum current injection that evoked at least a single AP. The inter-spike interval (ISI) was determined by finding the difference (in ms) between the crossing of 0 mV of one spike to the crossing of 0 mV by the next spike. To capture time-dependent changes in the frequency of APs, ISI 1/9 and 4/9 were determined by dividing the first ISI by the fourth and ninth ISI, respectively. ISI ratios were taken from the first sweep with at least 10 spikes. To determine the threshold voltage (V_thresh_) for an AP, for all spikes at each current injection, the derivative of the membrane voltage was taken across time to find the inflection point that corresponded with the beginning of the rising phase (i.e., threshold).

Threshold voltages for all spikes were then averaged to arrive at a single value of V_thresh_. Reset voltage (V_reset_) was defined as the minimum voltage value between spikes. A single value for V_reset_ was obtained in the same way as was done for V_thresh_. V_thresh_ and V_reset_ were used as parameters in the computational model.

#### Passive properties analysis

To obtain the R_in_, the difference between the baseline voltage (holding membrane voltage of approximately -65 mV) and the average voltage response to a - 20 pA injection (measured at steady state) was divided by that current injection value of 20 pA. The membrane time constant, 𝜏_m_, was the fitted response to the -20 pA injection. Membrane Capacitance (C_m_) was determined by dividing 𝜏_m_ by the R_in_. The resting membrane potential (V_m_) was measured as the potential before any current was injected. Finally, Voltage Sag Ratio was determined by dividing the difference between the minimum voltage at the peak deflection to a -500 pA current injection and the voltage of the steady state response by the difference between the minimum voltage at the peak deflection and the baseline voltage.

Electrophysiological data were analyzed using custom routines written in MATLAB, available at (https://github.com/CruzMartinLab/PV-mC4_Project/tree/main/ephys_code).

## Computational Model

### Model Neurons

Using the DynaSim toolbox (62), PYR and PV cells for each of the four networks – PV-mC4-WT and PV-mC4-KI male, and PV-mC4-WT and PV-mC4-KI female – were modeled as leaky integrate-and-fire neurons whose membrane voltage, V_m_, as a function of time, t, were given by the following:

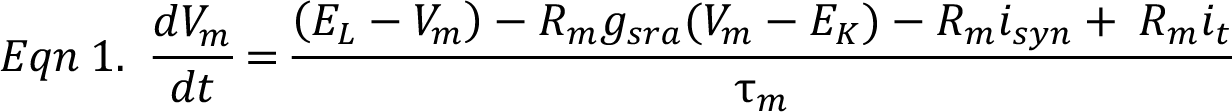

where E_L_ is the equilibrium potential, R_m_ is the membrane resistance, g_sra_ is the spike-rate adaptation, E_K_ is the potassium reversal potential, i_syn_ is the synaptic input, i_t_ is the applied current, and 𝜏_m_ is the product of R_m_ and membrane capacitance (C_m_). Values used for E_L_, R_m_, E_K_ (−102mV), and C_m_ were derived from the experimental data (**Table S5**), and differed from network to network only when there was a significant difference in the value of that variable between WT and KI mice overall or within-sex. If no significant difference was identified in a given parameter, the average across all sexes and conditions was taken and used uniformly across all networks.

### Establishing Model Neurons

For each of the four conditions, and by extension each of the four networks, a single model PYR and single model PV cell were generated. To create each model neuron, a waveform (i_t_) was matched to a subsection of the spiking protocol used during the experiment. The i_t_ waveform consisted of 17 sweeps of a square current pulse (500 ms) beginning at -10 pA, increasing in 30 pA steps, and finishing at 470 pA. From here, firing rates were extracted to construct frequency-current (FI) curves for each model cell that were used to compare to the experimental FI curves. An additional term, R_m_*i_std_, was added to the numerator of equation 1 (*Eqn 1*). This term served to shift the baseline voltage from V_m_ to -65 mV during the simulation, replicating the experimental conditions of current clamp under which the experimental data were acquired. i_t_ was multiplied by a scale factor, *I_act_*, representing active conductances and other intrinsic excitability properties which improved the fitting of the modeled FI curve to that of its experimental counterpart (**Fig. S5**). This same scale factor was used for i_syn_ to model the changes in intrinsic excitability. Reset voltage (V_reset_) and threshold voltages (V_thresh_) were determined experimentally, and the selection of the specific values used in each model cell followed the same logic as that for E_L_, R_m_, and C_m_. An increase in V_m_ above V_thresh_ constituted a spike, and V_m_ was then reduced to V_reset_ for an absolute refractory period, t_ref_, of 1 ms.

For all PYR and PV cells, spike rate adaptation time constants (𝜏_sra_) were set to 100 and 5 ms, respectively, reflecting the strong adaptation observed in regular-spiking PYRs and nominal levels of adaptation in fast-spiking PV cells (63–65). Implementation of spike-rate adaptation was accomplished by the following (66):

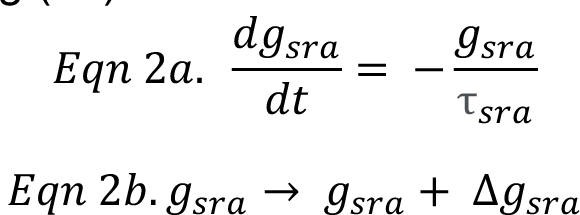

Moreover, spike-rate adaptation conductance was increased by an increment, Δg_sra_, which returns to zero in time 𝜏_sra_ (67). All model cells closely matched their experimental counterparts (**Fig. S5**).

### Establishing Networks

#### Simulating Network Noise

All PYR and PV model cells received random input to simulate network noise at levels that caused these model cells to fire at typical spontaneous rates observed experimentally, 3-4 Hz for PYRs and 28-35 Hz for PV cells (68,69). For each simulation, a random number of brief current pulses within a specified range to evoke the aforementioned basal firing rates was selected for PYR and PV model cells. Moreover, for each simulation, the time at which these pulses were applied to each cell were randomized as well.

#### Network Architecture

Individual networks were built to simulate WT male, KI male, WT female, and KI female conditions. For each of the four networks, fifteen 3 s simulations were run, the simulations serving as our statistical replicates.

Each network consisted of a PYR and PV circuit in a generalized input and output layer. In the input layer, PYR1 and PV1 simultaneously received an identical applied current, i_t._ In all network simulations, i_t_ consisted of a 4 Hz, fixed-amplitude sine wave atop the final 2 s of a 2.5 s, 50 pA DC component. The use of a non-varying DC component combined with a phasic sine wave input was used here to model dynamic input in an interpretable way (70,71) while preserving biological relevance (72–75) in our model. The first and final 250 ms of each simulation had no applied current. For all data plotted, only the 2 s in which the sine wave was applied was considered for analysis. Each of the fifteen simulations for a given network contained three trials in which the only variable altered was the peak of the sine wave (i.e., the peak of the applied current, i_t_): 200, 275, and 350 pA.

With respect to network connectivity, PYR1 synapses onto both PYR2 and PV2, in the output layer. PV1 synapses onto PV2. PV2 synapses onto PYR2. This specific feed-forward arrangement was selected such that all four basic connectivity sub-types (E->E, E->I, I->E, and I->I) sampled experimentally via mEPSCs and mIPSCs recorded in both PYR and PV cells could be included and thus be leveraged to better understand the effects of increased levels of *mC4* in PV cells on the firing properties of model neurons. This feed-forward network architecture is an adaptation of that used by Nocon et al. (76), but was further modified using arrangement principles similar to those used by Seay et al. (77) and Moore et al. (78). Notably, our model lacks feed-back circuitry and classical inhibitory action of other interneuron types, and favors single synapses over a population construction. Motivating these explicit simplifications for our current model was the benefit in interpreting the changes in the output model cells and an acknowledgement that the experimental data upon which many aspects of the model are founded come from *ex vivo*, rather than *in vivo*, data.

#### Synaptic Connectivity

To model postsynaptic currents (PSCs), after first setting all synaptic weights to the same fixed value across all four networks, weights were altered by adjusting the conductance of the inputs only where there were significant changes in the experimental mEPSC and/or mIPSC data recorded in PYR and PV cells (**Table S6**). These weights were adjusted by the same percent change as that observed in the mEPSC or mIPSC frequency or amplitude of the experimental data. In line with previous models (67,76,77), short-term depression was employed in all synapses. PSCs were modeled via the difference of two decaying exponential functions (76) featuring time constants where 𝜏_1_>𝜏_2_:

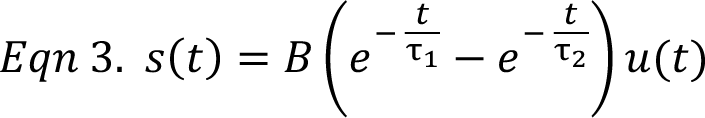

where B is a normalization constant, s = 1 at maximum, and u(t) is the unit step. From here, s can be represented by the following two ordinary differential equations:

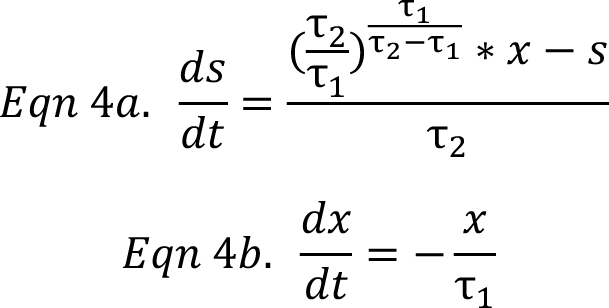

When a presynaptic spike is detected by a postsynaptic cell, x (spike inputs to that postsynaptic cell) is increased by P – representing synaptic strength – which is then reduced by f_p_, a fraction of its value (67,76).

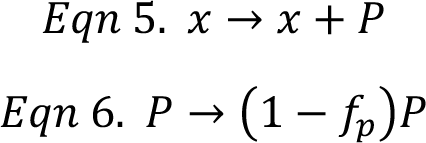

Recovery of P to 1 is defined by the time constant, 𝜏_p_, where:

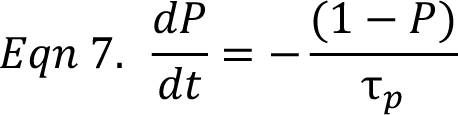

PSC rise (𝜏_1_) and decay (𝜏_2_) kinetics were derived from experimental values (**Table S6**). Values used for f_p_ and 𝜏_p_ were the same as those in Nocon et al. (76). In total, PSCs were modeled as:

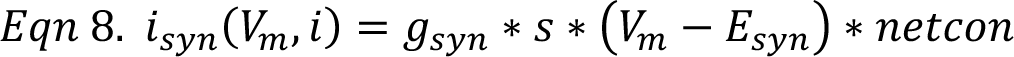

where g_syn_ is synaptic conductance, s is the time-dependent PSC defined above, E_syn_ is the reversal potential of the synaptic conductances, and netcon is a binary connectivity matrix in which rows represent sources and columns represent targets.

#### Additional Network Noise

An increase in network-level noise in response to suppression of PV cells has been directly demonstrated by multiple groups (5,79,80). After using our model to demonstrate PV cell hypoactivity in the male KI network, the basal network noise (3-4 Hz) delivered to PYR2 exclusively was increased by a factor of 1.5x (76). This increase in mean firing rate of PYR2 – representing an increase in network-noise in response to PV cell hypoactivity – has been used in previous models (76) and was set in order to match *in vivo* experimentally-derived changes in basal firing rates in response to PV suppression (5).

### Analysis of Network Parameters

For each of the four networks and the three peak values of the applied current, analysis was performed on 15 independent simulations.

#### Firing rate (FR)

Firing rate was calculated as the average number of spikes per second (Hz) during the 2 s in which the sine wave stimulus of the applied current occurred.

#### Transfer Entropy (TE)

TE is a metric used in determining connectivity in complex networks, considered to describe the effective flow of information between neurons (81,82). To determine the dependency of the spike train of PYR2 or PV2 on the spike train of PYR1 more directly, we calculated the TE (83) using a MATLAB Toolbox (81).

#### PYR1xPV2 Latency

Custom scripts written in MATLAB were used to determine the PYR1xPV2 latency. For each simulation, for each spike occurring in PYR1, the time (in ms) to the nearest spike in PV2 that followed the spike in PYR1 was calculated. The mean latency and latency standard deviation were calculated by averaging- or taking the standard deviation of, respectively, all latencies for a given simulation.

### Statistical Analysis

All statistical analyses were completed in GraphPad Prism 8.0, and the threshold for significance for all tests was set to 0.05 (α = 0.05). Full statistical reports for all plots are available in the Statistical Supplement document. Briefly, all behavior and electrophysiology plots representing sex-pooled data (WT males and females as one group being compared to KI males and females together as a second group) were analyzed by t-tests, t-tests with Welch’s correction, or Mann-Whitney tests. Moreover, all behavior and electrophysiology plots representing sex-separated data were analyzed by Two-way ANOVA’s with a Šídák’s multiple comparisons test only executed when appropriate. Cumulative frequency plots were analyzed with a Kolmogorov-Smirnov (KS) test. Frequency-current (FI) curves were analyzed using Repeated Measure Two-way ANOVAs. PV cell density was analyzed using a Mann-Whitney test. MFISH data were analyzed by Mann-Whitney tests or Two-way ANOVAs with Šídák’s multiple comparisons. Computational data were analyzed by Two-way ANOVAs with Šídák’s multiple comparison tests (firing rate and latency plots) and by Repeated Measure Two-way ANOVAs (transfer entropy plots). For all statistical analyses relating to the computational model, any *p*<0.05 was denoted in the figure by a ‘#’ sign – exact p-values can be found in the Statistical Supplement document. Figures were prepared using CorelDRAW Graphics Suite X8 (Corel Corporation) and ImageJ (NIH). Data are presented as the mean ± SEM, unless otherwise noted.

## RESULTS

### A novel transgenic mouse line permits PV cell specific overexpression of complement component 4

We generated a tunable conditional transgenic mouse based on a design by Dolatshad et al. (51) to reliably drive overexpression (OE) of *mC4* (mC4-OE) in specific cell types, and to incorporate genetic recombination switches that allow the conversion of different OE alleles for tunable transgene expression (**Fig. 1A**). Comer et al. (34) showed that mPFC neurons in postnatal day (P) 30 control mice express low levels of *C4b* transcript, which were not present in tissue from *C4b* knock-out mice (84).

**Figure 1.**
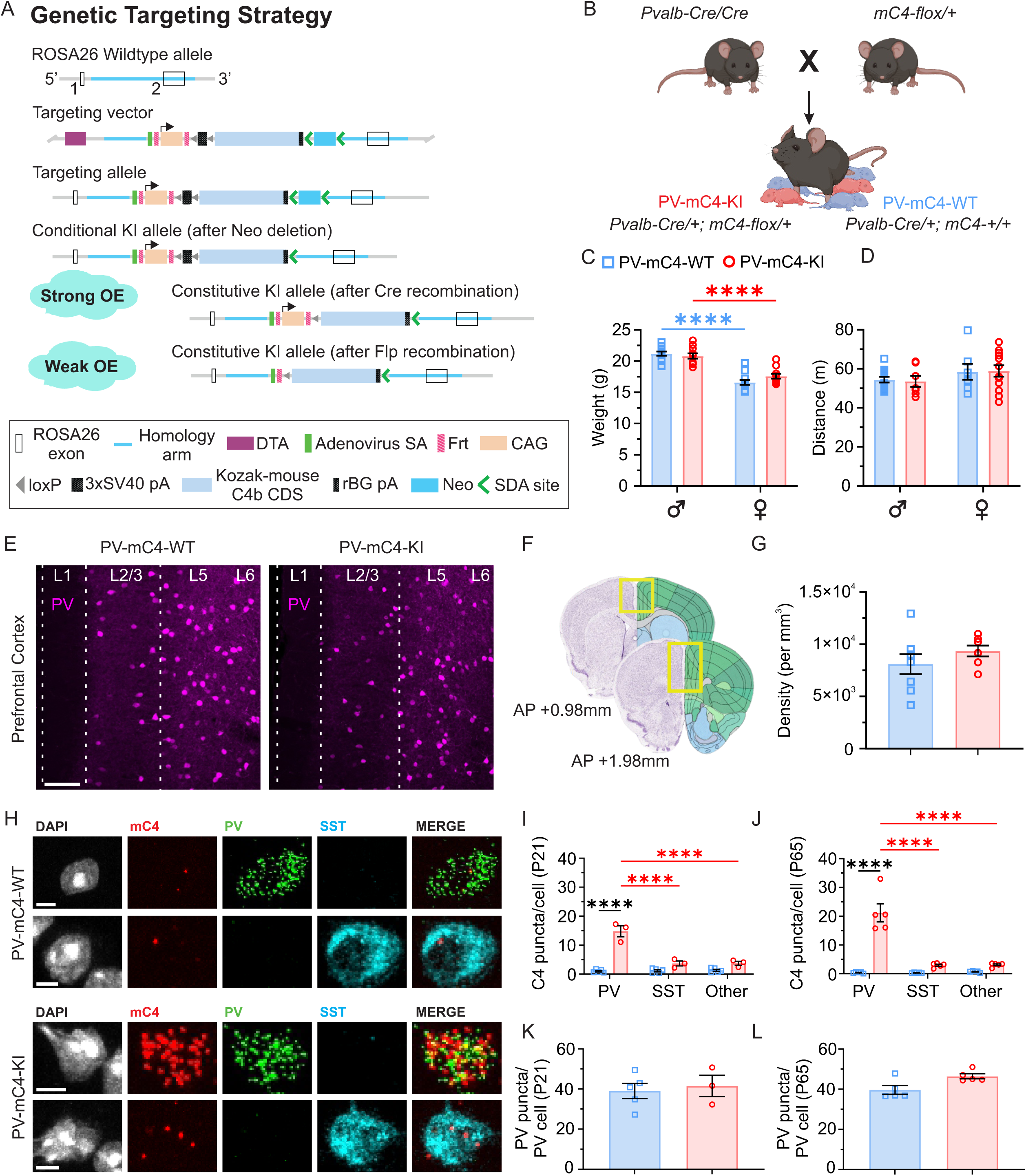
A novel transgenic mouse line permits PV cell specific overexpression of complement component 4. (**A**) Genetic strategy. Positive selection marker (Neo), self-deletion anchor (SDA) sites. Diphtheria toxin A (DTA) sites. The constitutive KI allele can be obtained after Cre- or Flp-mediated recombination, yielding relatively strong or weak OE of *mC4*, respectively. (**B**) Breeding scheme. (**C**) Increased levels of *mC4* in PV cells did not alter mouse weights compared to controls. (**D**) OE of *mC4* in PV cells did not impact the distance traveled. (**E**) Representative confocal images of PV cells (*magenta*) in mouse mPFC in P55-74 WT (*left*) and KI (*right*) animals. *scale bar* = 100 um. L, layer. (**F**) mPFC Cell density, yellow box. Bregma coordinates. AP, Anterior-posterior axis. (**G**) No difference in PV cell density between WT and KI. (**H**, *top two rows*) Representative confocal images (40x) of a PV (first row) and somatostatin (SST) cell (second row) in a WT mouse. (**H**, *bottom two rows*) Representative confocal images (40x) of a PV (first row) and SST cell (second row) in a KI mouse. For all rows in H, *left to right*: DAPI (*gray*), *mC4* (*red*), PV (*green*), SST (*cyan*), merged image without DAPI. (**I**) In P21 KI mice, *mC4* was overexpressed in PV, but not in SST or all other DAPI-labeled cells. In P21 KI mice, *mC4* expression was greater in PV than in SST and Other cells. (**J**) In P65 mice, *mC4* was overexpressed in PV, but not in SST or all other DAPI-labeled cells. In KI mice, *mC4* expression was greater in PV than in SST and Other cells. (**K**, **L**) No differences in PV expression between groups. WT: blue squares, KI: red circles. All statistics,\**p<*0.05, \*\**p* <0.01, \*\*\**p* <0.001, \*\*\*\**p* <0.0001. For information on statistics, see Statistical Supplement. Mean ± SEM shown.

To achieve specific OE of *mC4* in parvalbumin (PV)-positive cells (PV-mC4-OE) under the strong CAG promoter (85,86), we crossed PV-Cre mice with the conditional mC4-OE KI mouse line, mC4-KI mice (**Fig. 1A, B**). We generated litters that consisted of a mixture of pups that either inherited the floxed mC4-KI allele and thus overexpressed *mC4* specifically in PV cells (PV-mC4-KI, or KI) or did not inherit the floxed mC4-KI allele and were used as a wild type littermate control (PV-mC4-WT, or WT) (**Fig. 1B**). To control for effects of Cre-recombinase expression (87,88), we crossed homozygous PV-Cre mice to heterozygous mC4-KIs to obtain Cre recombinase expression in all offspring (**Fig. 1B**). We focused on a P40-60 temporal window, roughly equivalent to the young-adult life stage that immediately precedes SCZ symptom onset (89). PV-mC4-OE did not lead to observable health or locomotor deficits, as measured via weight (**Fig. 1C**) and distance traveled (**Fig. 1D**). Moreover, the density of PV cells was not significantly changed in PV-mC4-KI mice, compared to controls (**Fig. 1E-G**).

Next, we used multiplex fluorescence *in situ* hybridization (M-FISH) in the mPFC of WT and KI mice (**Fig. 1H-J**). At P21, we observed a significant increase in the number of *mC4* mRNA puncta in PV cells in PV-mC4-KI mice, indicating reliable mC4-OE in this interneuron type (**Fig. 1I**). We did not observe an increase in *mC4* puncta in neighboring somatostatin (SST)-positive cells and non-PV-non-SST-expressing cells (“Other”), suggesting that our genetic approach led to specific mC4*-*OE in PV cells (**Fig. 1I**). In PV-mC4-KI mice, the number of *mC4* puncta in PV cells was not different between P21 and 65 (**Fig. 1I, J**), indicating that we achieved stable OE across development. Additionally, other cell types did not show differences in *mC4* expression compared to WT mice, indicating cell-type-specific maintenance of the OE into adulthood (**Fig. 1I, J**). Finally, quantification of *Pv* mRNA puncta revealed that PV-mC4-OE did not drive any significant changes in the number of PV puncta per cell at either P21 (**Fig. 1K**) or P65 (**Fig. 1L**), suggesting that *Pv* expression was not altered by increased levels of *mC4* in this cell type.

In summary, we have developed and validated a novel conditional mouse model that, when combined with cre-driver lines, facilitates the study of distinct cellular sources of *mC4* overexpression. These results also indicate that we can reliably and specifically overexpress *mC4* in PV cells and that transgenic mice are devoid of gross motor or health defects.

### PV-specific mC4-OE causes an increase in anxiety-like behavior in male mice

Anxiety and mood disorders are highly prevalent in SCZ patients, manifesting during the early stages of the illness and prior to episodes of psychosis (90–93). To determine if PV-mC4-OE drives changes in anxiety-like behavior in P40-60 young adult mice, we used a behavioral battery, yielding a robust and reliable measure of behavior (60) (**Fig. 2A**). To assess arousal levels, we measured the time spent by mice in a field’s anxiogenic regions, namely the open or lighted areas (**Fig. 2B**).

**Figure 2.**
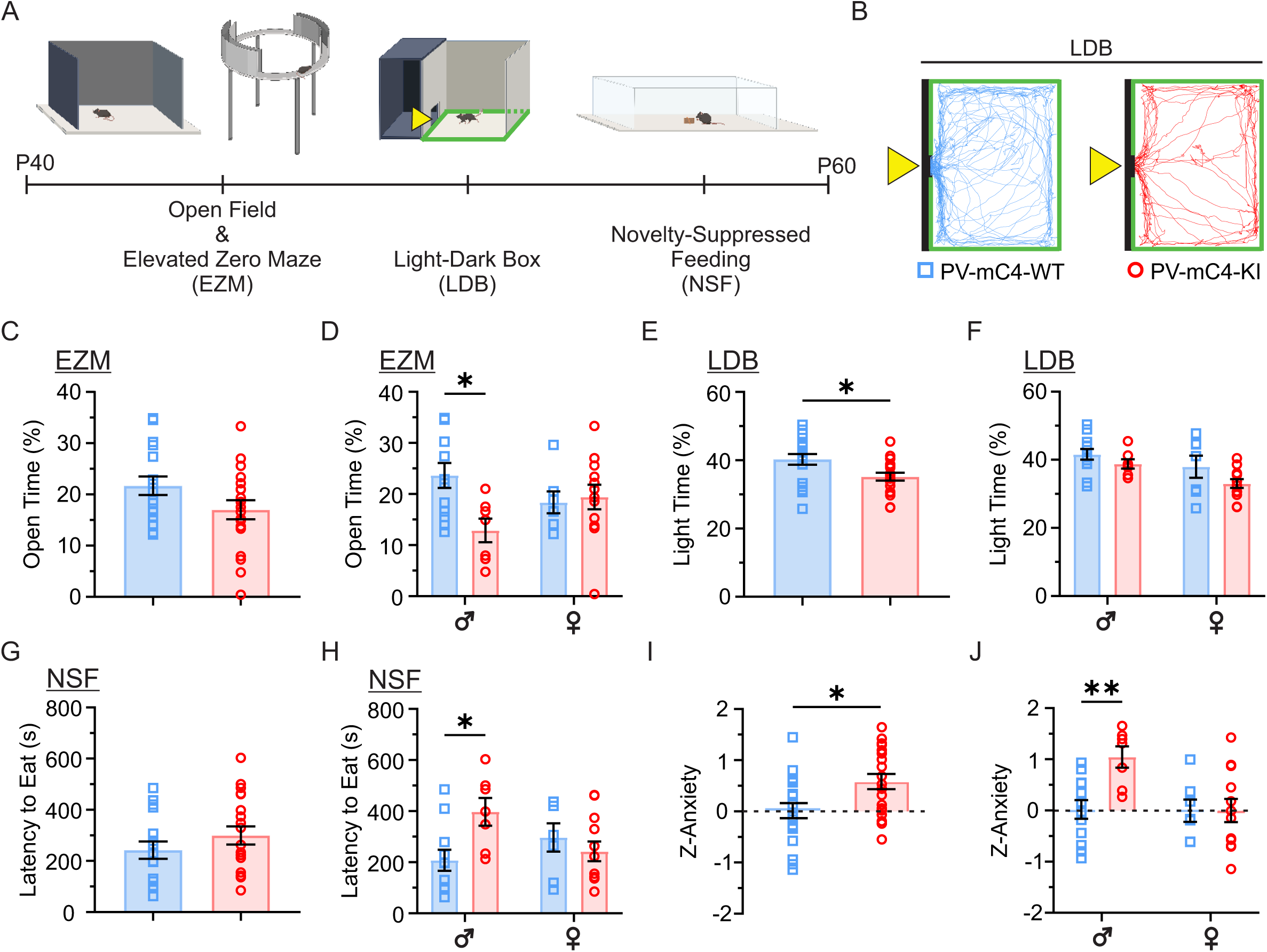
PV-specific mC4-OE causes an increase in anxiety-like behavior in male mice. (**A**) Timeline of behavioral assays. Mice were tracked in the light zone, represented by a green outline (B). (**B**) Representative trajectories of WT (*blue*) and KI *(red*) mice in the light zone of the LDB (colored traces). Yellow triangle, entrance to the light zone. (**C**) Percent time spent in the open arms of the EZM did not differ between groups. (**D**) Decreased time in the EZM open arms in male but not female KIs relative to WTs. (**E**) KI mice spent less time in the light zone of the LDB compared to WT mice. (**F**) No sex-dependent differences in LDB light zone time between groups. (**G**) Latency to feed in the NSF did not differ between groups. (**H**) Increase in the latency to feed for KI males compared to WT controls. (**I**) Significant increase in the Z-Anxiety of KI mice relative to WTs. (**J**) Compared to WT controls, there was an increase in the Z-Anxiety of KI male but not KI female mice. WT: blue squares, KI: red circles. All statistics,\**p*<0.05, \*\**p*<0.01. For information on statistics, see Statistical Supplement. Mean ± SEM shown.

In the Elevated Zero Maze (EZM), we observed no differences in the amount of time spent in the open arms between groups, suggesting that mice with increased *mC4* expression in PV neurons did not display an overall change in anxiety-like behavior, relative to WT controls (**Fig. 2C**). However, when we separated these data by sex, we observed a 46% reduction in the amount of time that KI males spent in the open arms, suggestive of a sexually dimorphic anxiety-like deficit (**Fig. 2D**). In contrast, we did not see such change in the behavior of KI female mice, relative to their WT controls (**Fig. 2D**). In the LDB, we observed a significant reduction in the time the PV-mC4-KIs spent in the light zone, again indicative of an increased anxiety-like response (**Fig. 2E**). Separation of the data by sex did not reveal the same sexual dimorphism as observed in the EZM (**Fig. 2F**). Lastly, in the novelty-suppressed feeding assay (NSF), we observed no change in the latency to feed between groups (**Fig. 2G**). Despite this, much like the EZM, there was a significant increase in anxiety-like behavior in the KI male mice, observed via a 91% increase in the latency to feed, but no such changes were observed in the PV-mC4-KI females (**Fig. 2H**). We performed the NSF in a safer environment, a mouse cage, where mice are exposed to bedding and can navigate close to the cage wall. In the Cage NSF, we did not observe differences in the latency to feed between groups (**Fig. S1**), suggesting that the NSF in the novel environment is anxiety-inducing and that the male KI mice, indeed, have increased anxiety-like behavior levels.

To describe the anxiety-like behavior of mice in response to PV-mC4-OE, we used a z-scoring approach that effectively tracks the performance of each mouse across all three behavioral assays, yielding a single score for each mouse, termed its Z-Anxiety score (60). We observed that the Z-Anxiety score of KI mice was increased in comparison to the control group (**Fig. 2I**). However, we discovered that overall heightened anxiety-like behavior is solely driven by a significant increase in the Z-Anxiety of PV-mC4-KI male mice (**Fig. 2J**). In support of this, female KI mice did not exhibit an increased Z-Anxiety score compared to the WT controls (**Fig. 2J**). In summary, PV-mC4-OE causes sex-dependent behavioral changes, with male KI mice exhibiting increased anxiety-like behavior relative to WT controls.

### Increased levels of *mC4* in PV cells disrupts active but not passive social behaviors

We employed a naturalistic, freely-moving interaction assay between experimental mice and a novel, juvenile, sex-matched CD-1 stimulus mouse to determine whether increased levels of *mC4* in fast-spiking cells led to social behavioral changes. (**Fig. 3**). First, both male and female KI mice spent a similar time interacting with a novel object as their WT counterparts (**Fig. 3A, B**), suggesting that PV-mC4-OE does not alter novelty-seeking behaviors or general motivation.

**Figure 3.**
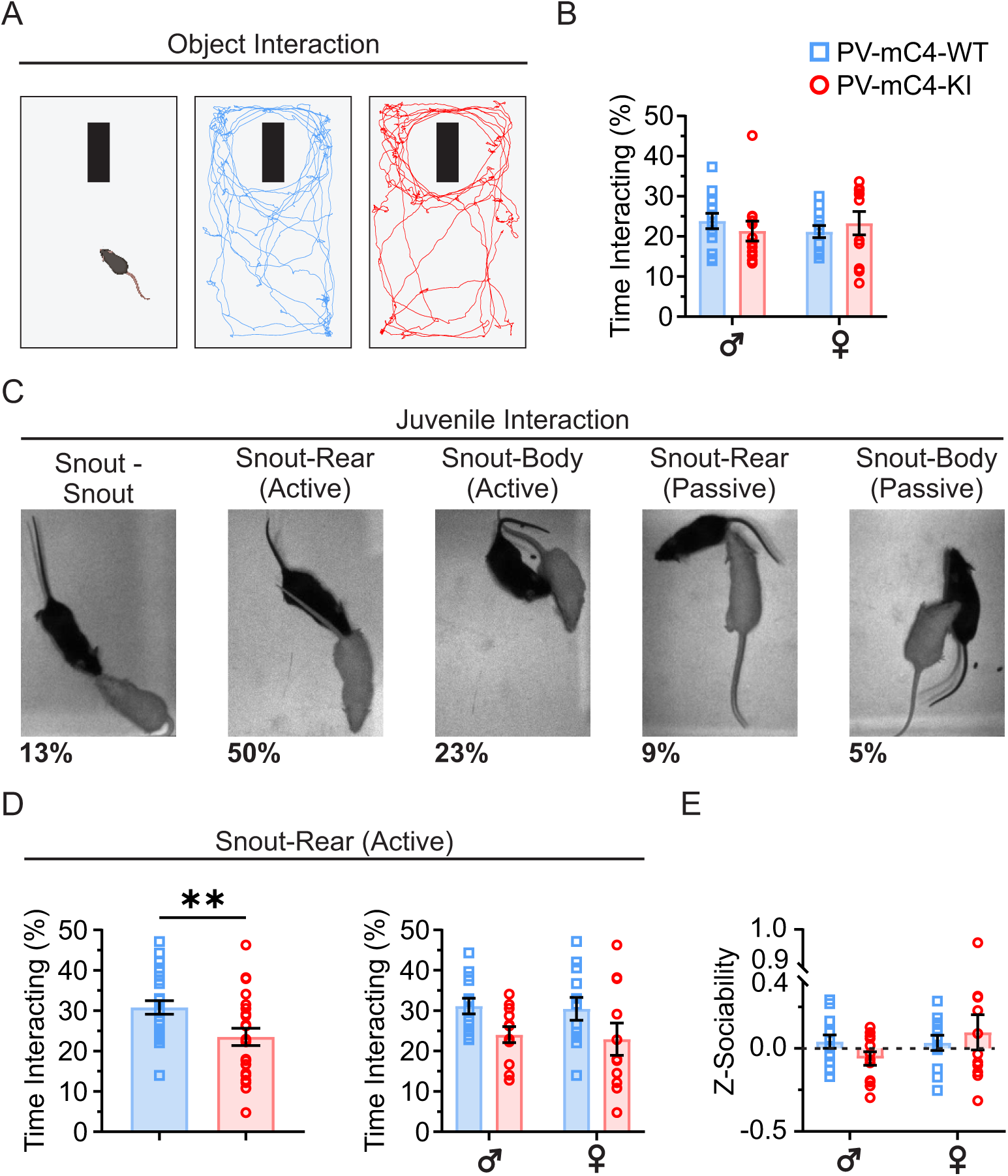
Increased levels of *mC4* in PV cells disrupts active but not passive social behaviors. (**A**) Object interaction task (*left*). Representative trajectories of WT (*middle, blue*) and KI (*right, red*) exploring a novel object (*black rectangle*). (**B**) KI mice explored the novel object as much as WT controls. (**C**) Representative images of interaction sub-classes. *Active*, experimental mouse engaging the stimulus mouse; *Passive*, stimulus mouse engaging the experimental mouse. Percentage of each sub-class of behavior is below. (**D**) *Left*: Relative to WT controls, KI mice spent less time engaged in the active snout-rear interaction type. *Right*: There were no sex-related differences in active-snout-rear interaction between groups. (**E**) There was no change in Z-Sociability in either KI male or female mice. WT: blue squares. KI: red circles. All statistics,\**p*<0.05, \*\**p*<0.01. For information on statistics, see Statistical Supplement. Mean ± SEM shown.

Immediately following the object interaction assay, the object was replaced with a stimulus mouse (**Fig. 3C**). Interactions between WT or KI mice and the stimulus mouse were divided into one of five sub-classes (**Fig. 3C**). Of all interaction sub-classes, experimental mice engaged in the active (active: experimental mouse initiating the interaction vs. passive: stimulus mouse initiating the interaction) snout-rear interaction (snout-ano-genital interactions) the most frequently, comprising half of all interaction time across all groups (**Fig. 3C**, *percentages*).

Notably, PV-mC4-KI mice engaged less in active interactions than controls (**Fig. S2A**); this reduction was driven by a 24% decrease in active snout-rear interaction (**Fig. 3D**). These results indicate that increased levels of *mC4* in PV cells lead to a reduction in snout-ano-genital interactions. Unlike the dimorphic nature of the anxiety-like behavior, deficits in sociability affected KI mice of both sexes (**Fig. 3D**).

We observed more active than passive interactions by the experimental mice, which suggests that the experimental mice initiated more social interactions than the stimulus mice. (**Fig. 3C**, **Fig. S2A**). Furthermore, we did not notice any changes in passive interactions, indicating that stimulus mice behaved similarly when interacting with both groups (**Fig. S2**). In the remaining interaction sub-classes, which were less frequent than the active snout-rear interaction, WT and KI mice interacted similarly with the stimulus mouse (**Fig. S2B-I**).

Computing a Z-Sociability score that accounts for all five interaction sub-classes did not reveal a broad deficit in sociability (**Fig. 3E**). Separately, in a three-chamber assay of sociability, a more restricted paradigm, PV-mC4-OE did not alter the social preference of PV-mC4-KIs, relative to WTs (**Fig. S3**), suggesting all experimental animals preferred to interact with a mouse rather than an object. Our results also suggest that the freely-moving interaction assay allowed us to capture the complex behavioral phenotype that OE mice exhibit.

Although female PV-mC4-KI mice did not exhibit changes in anxiety-like behavior, they had decreased social interactions as part of the KI group (**Fig. 3D**), indicating that the mechanisms driving the pathology in the anxiety-like and social behaviors in response to PV-mC4-OE likely function independently. Overall, these results suggest that increased levels of *mC4* in PV cells disrupt the circuits that underlie emotional and social behavior in mice.

### Sex-related differences in excitatory-inhibitory dynamics in mPFC PV cells with increased levels of *mC4* in PV cells

To determine whether PV-mC4-OE changed the connectivity of circuits in the mPFC, we performed whole-cell voltage-clamp recordings in acute brain slices. Specifically, we first recorded miniature excitatory post-synaptic currents (mEPSCs) in PV neurons in layer (L) 2/3 of the mPFC in P40-60 mice (**Fig. 4A-G**), the same temporal window within which we identified behavioral deficits in response to PV-mC4-OE. Though we observed no change in PV cell mEPSC amplitude in PV-mC4-KI mice (**Fig. 4A-C**), we observed a 39% reduction in mEPSC frequency specifically in KI male mice (**Fig. 4D**). In contrast, there were no changes in PV cell mEPSC frequency in the KI females (**Fig. 4D**). In support of this, there was a rightward shift in the frequency distribution of inter-event-intervals (IEIs) in PV neurons in PV-mC4-KI males, but not females (**Fig. 4E**), suggesting that PV-mC4-OE leads to a decrease in excitatory drive to this fast-spiking neuron. Finally, PV-mC4-OE did not alter the Rise_10-90_ or Decay_tau_ of mEPSCs recorded in PV cells (**Fig. 4F, G**), suggesting that increased levels of *mC4* did not alter the kinetics of the PV cell postsynaptic response.

**Figure 4.**
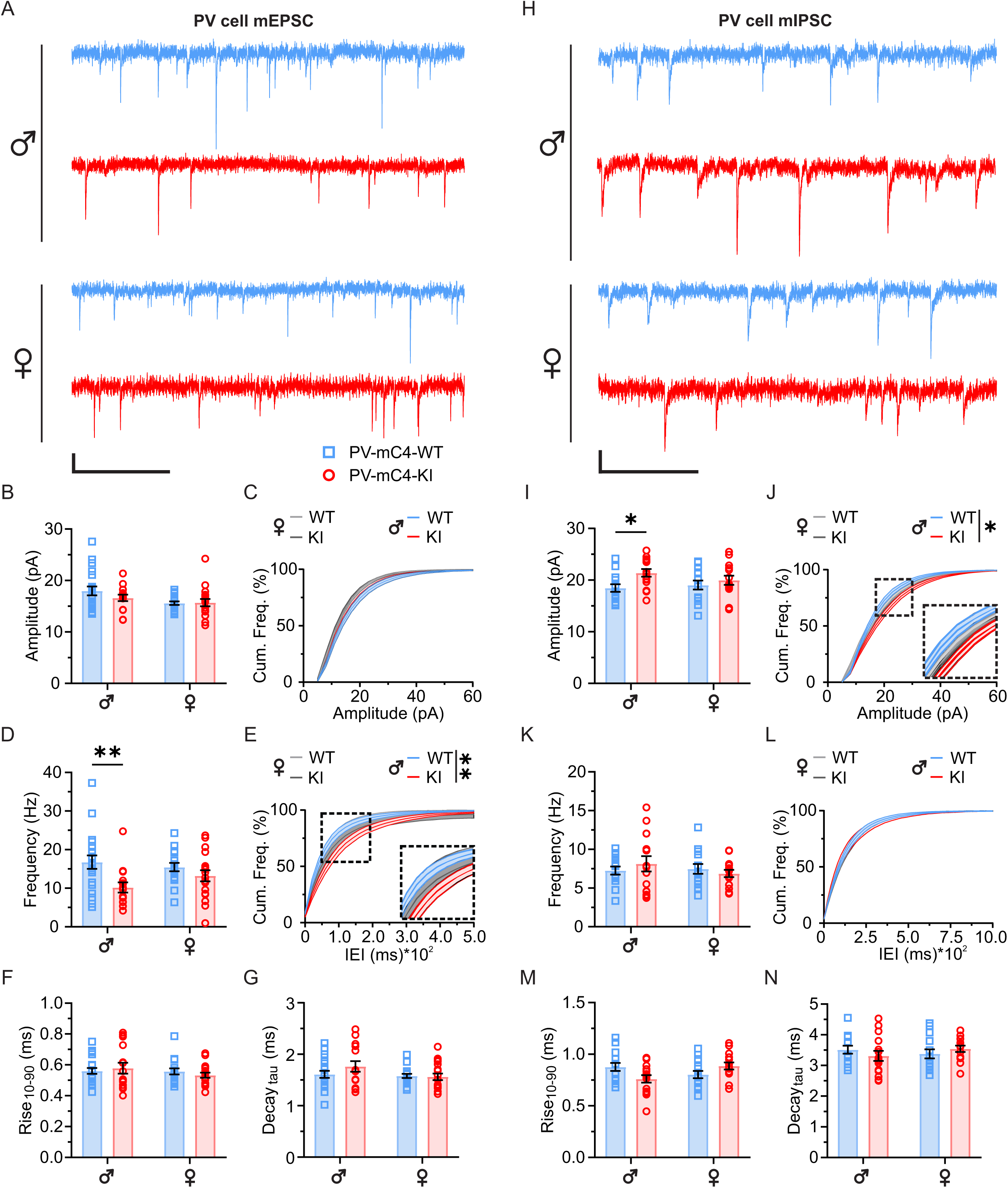
Sex-related difference in excitatory-inhibitory dynamics in mPFC PV cells with increased levels of *mC4* in PV cells. (**A**) Representative voltage-clamp traces of mEPSCs recorded in PV cells of P40-60 WT (*blue*) and KI (*red*) mice, *scale bar* = 250 ms, 10 pA. (**B**) No change in mEPSC amplitude in KI mice, compared to controls. (**C**) No shift in the cumulative frequency distribution of mEPSC amplitudes in KI mice. (**D**) mC4-OE led to a decrease in mEPSC frequency in KI male mice, but not KI female mice. (**E**) Increased *mC4* expression caused a rightward shift in the distribution of mEPSC inter-event-intervals (IEIs) of PV cells in KI male mice, but not KI female mice. (**F**) mEPSC Rise_10-90_ was not impacted in KI mice, relative to WT controls. (**G**) mEPSC Decay_tau_ was not changed in KI mice. (**H**) Representative 1s voltage clamp traces of mIPSCs recorded in PV cells of P40-60 WT (*blue*) and KI (*red*) mice, *scale bar* = 250 ms, 10 pA. (**I**) mIPSC amplitude was increased in KI male, but not KI female mice. (**J**) Increased expression of *mC4* caused a rightward shift in the distribution of mIPSC amplitudes in KI male mice. (**K**) mIPSC frequency was not changed in KI mice. (**L**) No shift in the distribution of mIPSC IEIs of KI mice. (**M**) Relative to controls, there were no changes in mIPSC Rise_10-90_ with increased levels of *mC4* in PV cells. (**N**) mIPSC Decay_tau_ was not changed in KI mice. WT: blue squares. KI: red circles. *N* represents cells. All statistics,\**p*<0.05, \*\**p*<0.01. For information on statistics, see Statistical Supplement. Mean ± SEM shown.

To determine if PV-mC4-OE impacted the inhibitory drive to PV neurons, we recorded miniature inhibitory post-synaptic currents (mIPSCs) in this inhibitory cell type (**Fig. 4H-N**). Using a high-chloride internal recording solution with a chloride reversal potential of -13 mV yielded inward mIPSCs when recording at -70 mV (**Fig. 4H**). The recordings revealed that PV-mC4-OE drove a 16% increase in the amplitude of PV cell mIPSCs (**Fig. 4I**) and a rightward shift in the distribution of mIPSC amplitudes (**Fig. 4J**) specifically in male, but not female mice, suggesting that inhibitory inputs are increased in PV cells in male PV-mC4-KI mice. Additionally, we observed no changes in mIPSC frequency (**Fig. 4K**, **L**), Rise_10-90_ (**Fig. 4M**), or Decay_tau_ (**Fig. 4N**) between groups.

Taken together, these results suggest that PV-mC4-OE drives sex-dependent alterations in PV cell excitatory and inhibitory connections, mirroring the sexually dimorphic changes in anxiety-like behavior. The combined effects of reduced excitation and increased inhibition to PV cells in KI male mice suggests hypoactivity of mPFC inhibitory circuits in response to increased levels of *mC4* in fast-spiking cells.

### PV-specific mC4-OE leads to opposing changes in the excitability of PV cells in male and female mice

We evaluated the passive and active properties of both mPFC L2/3 PV cells (**Fig. 5A-E**, **Tables S1, S2**) and PYRs (**Fig. 5F-J**, **Tables S3, S4**). To accomplish this, we injected steps of hyperpolarizing and depolarizing current pulses and recorded the membrane voltage (V_m_) changes. Changes in excitability of PV neurons in male and female mice in response to PV-mC4-OE diverged: while there was a significant decrease in PV cell spike frequency in KI male mice relative to WT males (**Fig. 5A**, **B**), we observed a significant increase in the spike frequency of PV cells in female OE mice, compared to WT females (**Fig. 5A**, **C**). This increase in excitability in PV cells in KI females was also accompanied by a 26% reduction in their rheobase, another indication of increased excitability (**Fig. 5D**). Finally, we observed a significant shift towards a more depolarized resting membrane voltage in KI mice overall (**Fig. 5E**, **Tables S1, S2**). These results suggest that there is a sex-dependent divergence in the fast-spiking cell’s excitability with higher *mC4* levels in PV cells.

**Figure 5.**
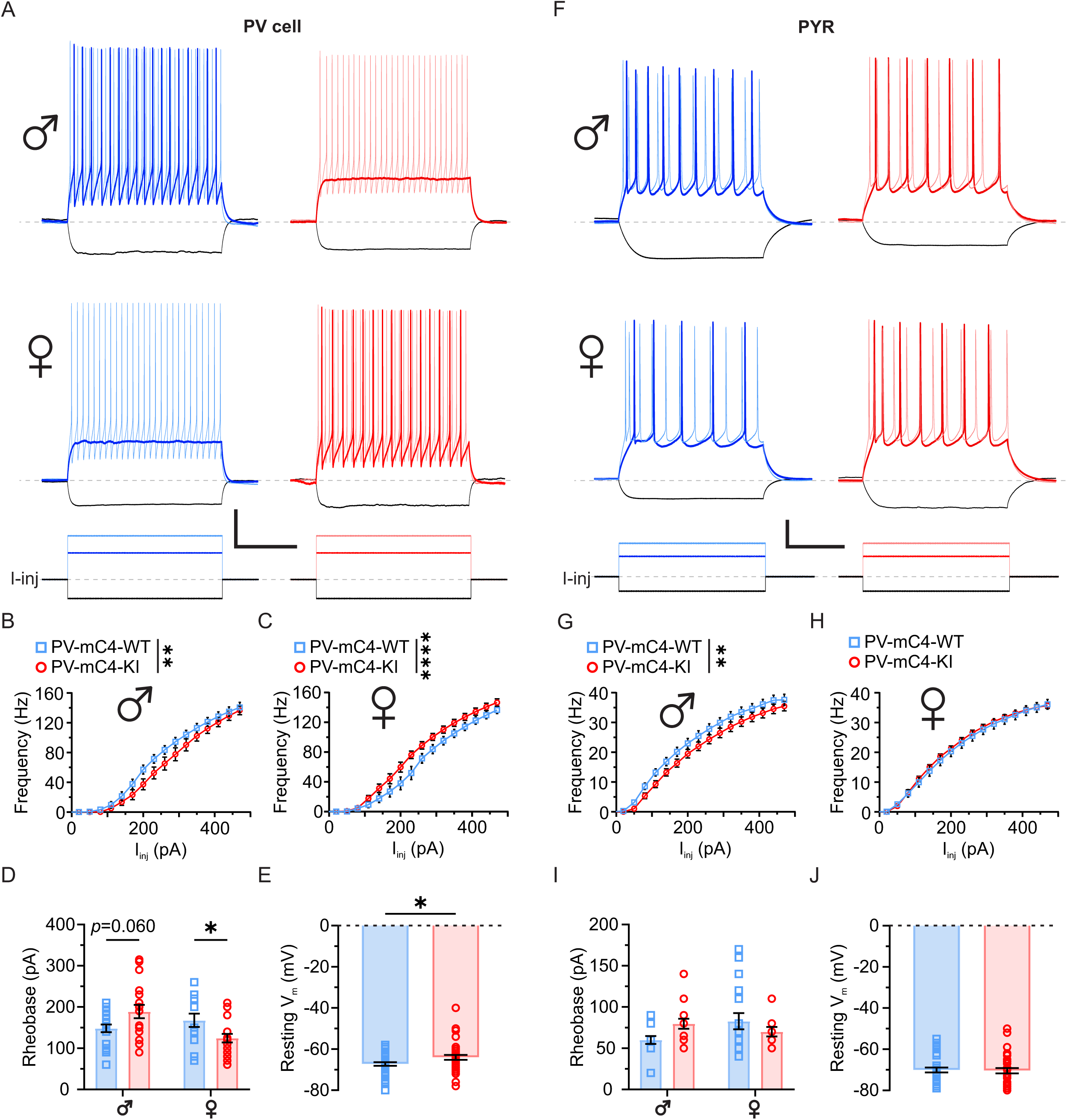
PV-specific mC4-OE leads to opposing changes in excitability of PV cells in male and female mice. (**A**) Representative recordings of PV cells injected with -100 (*black*), 140 (*darker colored shade*), and 230 pA (*lighter colored shade*) of current from WT (*blue*) and KI (*red*) mice, *scale bar* = 200 ms, 200 pA/20 mV. Baseline V_m_, -65 mV. (**B**) PV cells in KI male mice spike less than PV cells in WT male mice. (**C**) mC4-OE led to an increase in the excitability of PV cells in KI female mice, relative to controls. (**D**) Rheobase was decreased in KI female mice, relative to controls. (**E**) PV-mC4-OE drove a shift in PV cell resting membrane voltage towards a more depolarized V_m_. (**F**) Representative recordings of PYRs injected with -100 (*black*), 140 (*darker colored shade*), and 230 pA (*lighter colored shade*) from WT (*blue*) and KI (*red*) mice, *scale bar* = 200 ms, 200 pA/20 mV. (**G**) OE of *mC4* decreased the spike frequency of PYRs in KI male mice, relative to controls. (**H**) No differences in the excitability of PYRs in females between groups. (**I**) No changes in PYR Rheobase. (**J**) No change overall in PYR resting V_m_ in KI mice, compared to controls. WT: blue squares. KI: red circles. *N* represents cells. All statistics,\**p*<0.05, \*\**p*<0.01. For information on statistics, see Statistical Supplement. Mean ± SEM shown.

Though mC4-OE is limited to PV cells in this mouse model, it is possible that disruption in the activity of PV cells may elicit compensatory changes in PYRs. To this end, we recorded the membrane voltage response as before, now in mPFC L2/3 PYRs (**Fig. 5F-J**). Similar to PV neurons, in male mice PV-mC4-OE drove a reduction in the spike frequency of PYRs (**Fig. 5F**, **G**). Unlike PV cells, we observed no changes in spike frequency in the PYRs of KI female mice (**Fig. 5F**, **H**). Moreover, PV-mC4-OE did not alter the rheobase (**Fig. 5I**) or resting membrane voltage (**Fig. 5J**) of PYRs. These results indicate that PV-mC4-OE induced changes in PYR excitability in male mice.

Overall, increased *mC4* levels in PV cells caused sexually dimorphic effects on the excitability of cortical cells. Fast-spiking cells and PYRs in males showed a decrease in excitability, while fast-spiking cells in females exhibited hyperexcitability. This divergent outcome suggests that the male and female mouse brain respond to complement dysfunction in opposing ways.

### PV-specific mC4-OE alters the kinetics of mEPSCs in PYRs of female mice

We recorded mEPSCs (**Fig. S4A-E**) and mIPSCs (**Fig. S4F-J**) in L2/3 mPFC PYRs. PV-mC4-OE did not lead to changes in PYR mEPSC amplitude (**Fig. S4B**) or frequency (**Fig. S4C**) compared to controls. While the Rise_10-90_ of the mEPSCs in PYRs was also not altered in PV-mC4-KI mice (**Fig. S4D**), PV-mC4-OE caused a 14% reduction in Decay_tau_ of the mEPSCs in KI female, but not KI male mice (**Fig. S4E**), suggesting a change in receptor subunit composition in PYRs (94) or a change in the location of excitatory synapses along its somatodendritic axis (95,96). Finally, PV-mC4-OE did not induce changes in PYR mIPSC amplitude (**Fig. S4G**), frequency (**Fig. S4H**), or kinetics (**Fig. S4I**, **J**) relative to controls. These results suggest that PV-mC4-OE largely does not alter mPFC PYR synaptic drive.

### No changes in anxiety-like behavior with pan-neuronal overexpression of *mC4*

Next, we crossed the mC4-KI mouse to the BAF53b-Cre transgenic mouse line (50) that express Cre recombinase under the control of the mouse *Actl6b* gene promoter to drive mC4-OE in all neurons (PanN-mC4-OE). The expression of the BAF53b gene in neurons can first be detected during embryonic day 12.5 in the brain and spinal cord (97). Using a similar breeding strategy as with the PV-mC4-KI mice, litters consisted of a mix of mice that inherited the floxed mC4-KI allele and thus overexpressed *mC4* in all neurons (PanN-mC4-KI), or littermates that did not inherit the floxed mC4-KI allele, and thus were effectively WT (PanN-mC4-WT) (**Fig. 6A**, *top*).

**Figure 6.**
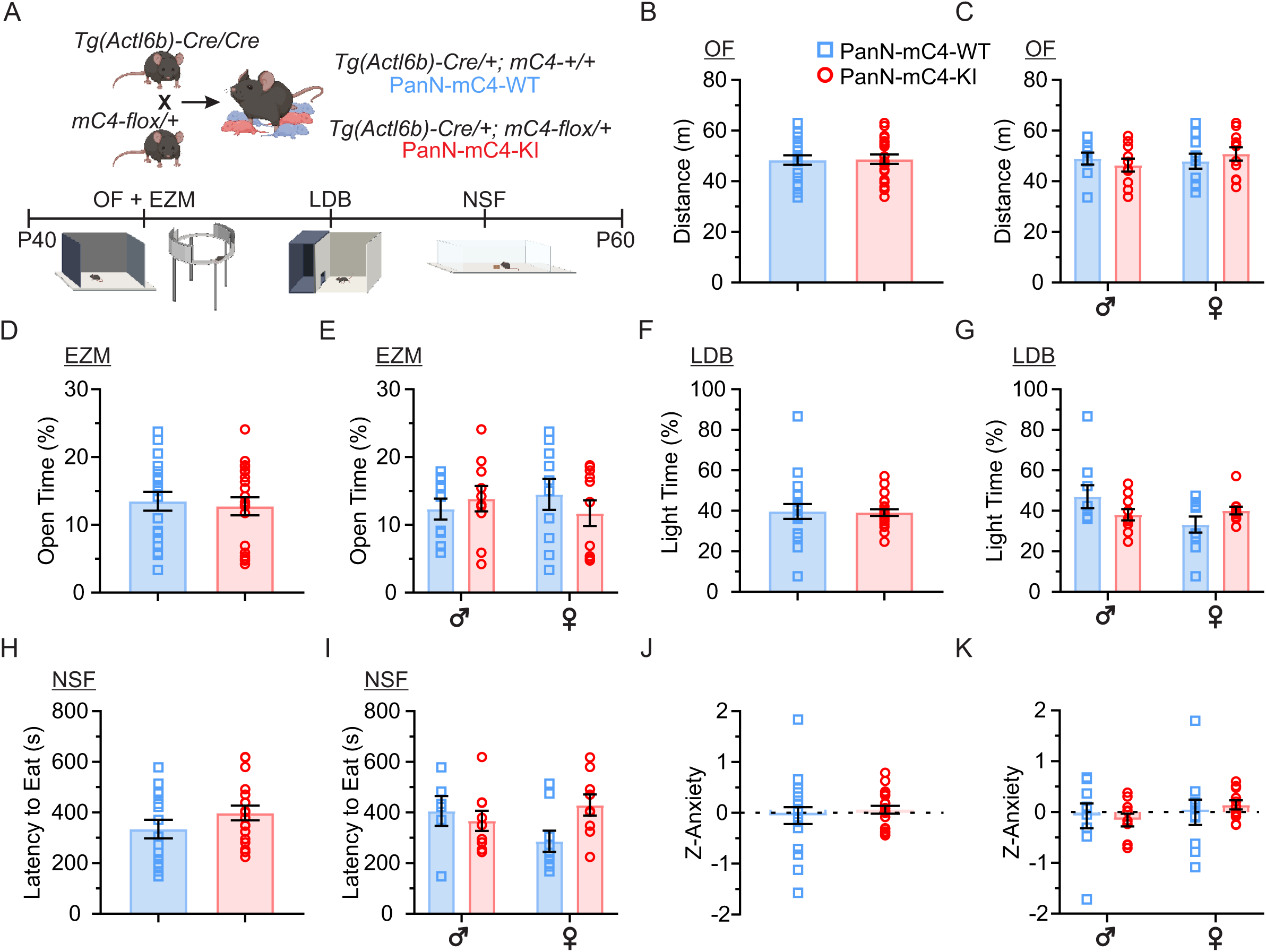
No changes in anxiety-like behavior with pan-neuronal overexpression of *mC4*. (**A**) Mouse breeding (*top*) and anxiety behavioral battery (*bottom*) as in Fig. 2. (**B**, **C**) Pan-neuronal mC4-OE does not alter locomotion. (**D**, **E**) No change in time spent in the open arms with pan-neuronal mC4-OE. (**F**, **G**) Time spent in the light zone was not altered by pan-neuronal mC4-OE. (**H**, **I**) OE of *mC4* in neurons did not alter feed latency. (**J**, **K**) mC4-OE in neurons did not lead to changes in overall anxiety-like behavior. WT: blue squares. KI: red circles. All statistics, \**p*<0.05. For information on statistics, see Statistical Supplement. Mean ± SEM shown.

We employed the same assays – the EZM, LDB, and NSF – to test anxiety-like behavior in P40-60 PanN-mC4-WT and PanN-mC4-KI mice (**Fig. 6A**, *bottom*). First, OE of *mC4* in all neurons did not alter the distance traveled compared to controls (**Fig. 6B**, **C**), suggesting that PanN-mC4-KI mice exhibited intact locomotion. Moreover, compared to controls, we did not observe any deficit in anxiety-like behavior in PanN-mC4-KI mice in the EZM (**Fig. 6D**, **E**), LDB (**Fig. 6F**, **G**), or NSF (**Fig. 6H**, **I**). Also, we did not observe any increase in Z-Anxiety in PanN-mC4-KI mice (**Fig. 6J**), nor in either sex specifically (**Fig. 6K**). Taken together, PanN-mC4-OE does not drive anxiety-like behavior in mice.

We have demonstrated that increased levels of *mC4* in PV neurons resulted in a strong, sex-specific anxiety-like phenotype not observed in pan-neuronal *mC4* overexpressors. This suggests that specific complement changes in PV cells leads to developmental dysfunction of inhibitory circuits that is more detrimental to brain function than pan-neuronal alterations.

### Disrupted neural communication and hyperexcitability in a network model of male mice with increased levels of *mC4* in PV cells

We used a computational model to determine how PV-mC4-driven deficits in connectivity and excitability of PYR and PV cells contribute to circuit-level abnormalities in a simulated network. Utilizing the DynaSim toolbox (62), we developed four networks with identical architecture (**Fig. 7A**) representing the experimental conditions – WT and KI male, and WT and KI female groups. The electrophysiological properties of individual PYR and PV cells – and their connectivity to one another – were matched to the experimental data (**Tables S5, S6**). Specifically, we first established unique models of PV and PYR units for each group using our experimental data (**Table S5**). To determine if these model units accurately reflected the experimental data, we compared the frequency vs. current (FI) curves of each model unit against its equivalent experimental cell (**Fig. S5**). In all cases, these model units accurately approximated their experimental counterpart.

**Figure 7.**
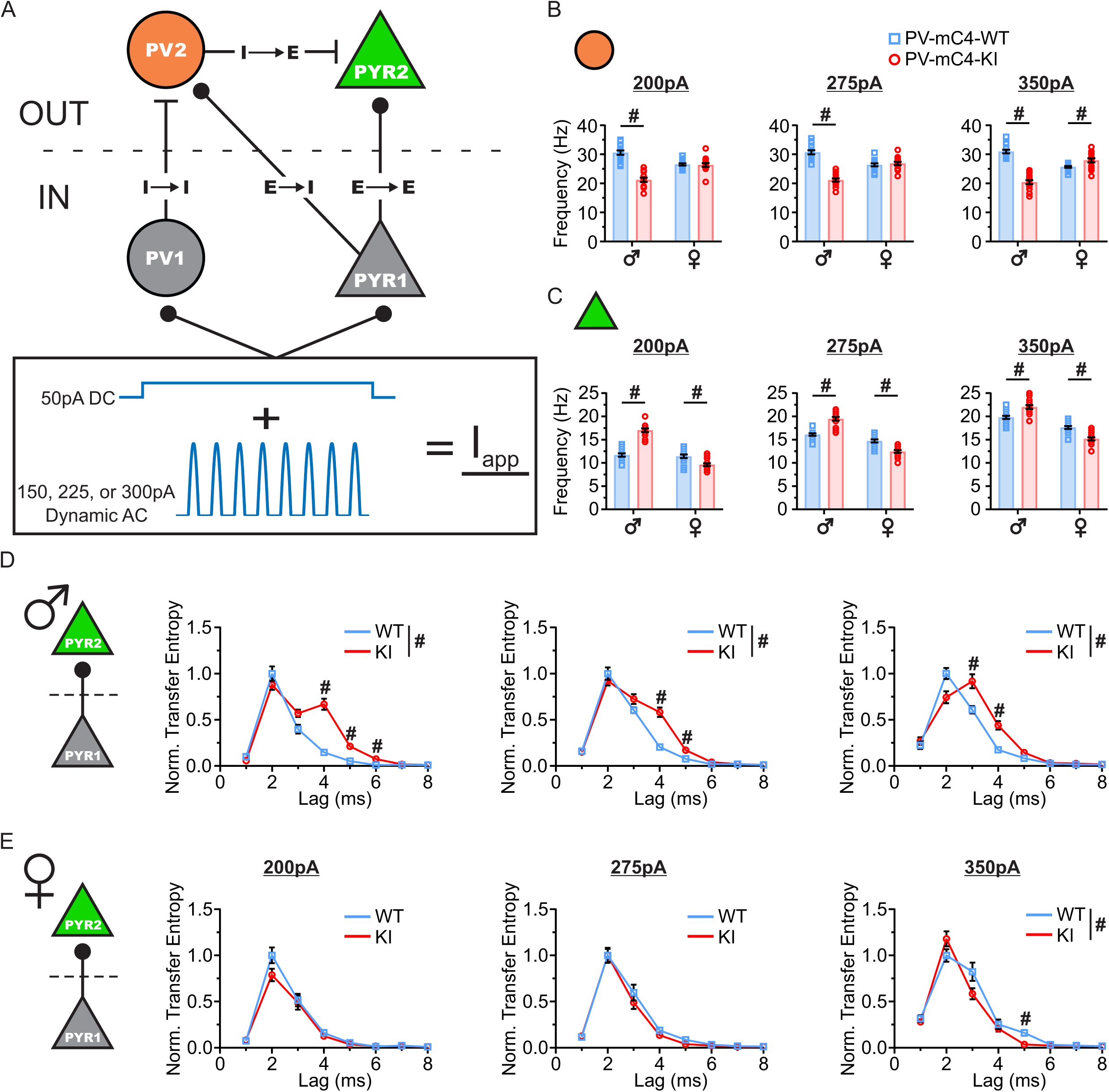
Disrupted neural communication and hyperexcitability in a network model of male mice with increased levels of *mC4* in PV cells. (**A**) Network architecture and applied current (*Iapp*). (**B**) The firing rate (FR) of PV2 in WT and KI networks at peak *I_app_* of 200 pA (*left*), 275 pA (*middle*), and 350 pA (*right*). (**C**) The FR of PYR2 in WT and KI networks at peak *I_app_* of 200 pA (*left*), 275 pA (*middle*), and 350 pA (*right*). (**D)** Transfer entropy (TE) at each delay (lag) of PYR1 onto PYR2 in male WT (*blue*) and male KI (*red*) networks at peak *I_app_* of 200 pA (*left*), 275 pA (*middle*), and 350 pA (*right*). (**E**) TE at each delay (lag) of PYR1 onto PYR2 in female WT (*blue*) and female KI (*red*) networks at peak *I_app_* of 200 pA (*left*), 275 pA (*middle*), and 350 pA (*right*). WT network: blue squares. KI network: red circles. *N*, simulations. *N*=15 for all networks. For information on statistics, see Statistical Supplement. Mean ± SEM shown. For all plots, ‘*#*’ indicates all *p*<0.05.

We hypothesized that downstream PYR in KI males would become hyperactive as a function of reduced inhibition. In support of this, in the output layer of the male KI neural network model we observed a significant decrease in the firing rate (FR) of PV2 (**Fig. 7B**) and a significant increase in PYR2 FR (**Fig. 7C**) at all three peak values of the applied current (I_app_), compared to the male WT network. This suggests that in the network model, increased levels of *mC4* in male PV model cells cause decreased activity of this fast-spiking model neuron, driving hyperactivity of PYR model cells.

To determine if the changes associated with PV-mC4-OE disrupt neuronal communication, we measured the transfer entropy (TE) of the direct PYR1->PV2 (**Fig. S6A-C**) and PYR1->PYR2 (**Fig. 7D, E**) connections. Specifically, we first investigated the likelihood that a spike in PYR1 would cause a spike in PV2 (**Fig. S6B, C**). In line with our hypothesis, we observed a significant reduction in the PYR1->PV2 TE in the male KI network at lag times of 1-3 ms at all peak values of I_app_ (**Fig. S6B**). This suggests that the effective transfer of information from PYR1 to PV2 is disrupted in the male KI network. This finding was further supported by a significant increase in both the average (**Fig. S6D**) and standard deviation (**Fig. S6E**) of the latency from any given PYR1 spike to the nearest following spike observed in PV2. Consistent with this compromised PYR1->PV2 communication, because PV2 is the only line of inhibition to PYR2 in the network, the PYR1->PYR2 TE is also significantly altered in the male KI network (**Fig. 7D**). Namely, we observed a broadening of the lag times over which activity in PYR1 could drive changes in PYR2 activity (**Fig. 7D**), indicating that the precise temporal relationship of PYR1->PYR2 communication is disrupted in the model male KI network compared to controls.

In KI female mice, we observed a significant increase in the intrinsic excitability of PV cells in response to PV-mC4-OE (**Fig. 5**). Interestingly, we observed no changes in the PV2 FR in the female KI network in either the 200 or 275 pA peak I_app_, compared to the female controls (**Fig. 7B***, left, middle*). However, when the peak of I_app_ was raised to 350 pA, we observed a significant 9% increase in the FR of PV2 in the female KI network (**Fig. 7B**, *right*), suggesting that at lower peak applied current values, the increase in intrinsic excitability of PV cells in the female KI network – and the potential increase in PV2 FR that may be expected in response to input from PYR1 – is neutralized by the increased spiking activity of PV1 (and thus PV1 inhibition to PV2). However, our results suggest that when we applied stronger stimulation, the intrinsic excitability of PV2 and its resulting increase in FR outweighs the influence of increased inhibition.

Notably, in the female KI network, despite observing no change in PV2 FR at peak I_app_ values of 200 and 275 pA, the FR of PYR2 is still significantly decreased compared to controls. Provided that PV2 is the only source of inhibition to PYR2, this finding of a lack of change in PV2 FR but a decrease in FR of PYR2 appears incongruous. To determine if it is changes not in the number of spikes, but in the timing of spikes of PV2 relative to PYR2 that may be driving this decrease in PYR2 FR in the female KI network, we compared the PYR1->PV2 TE in the female WT and KI networks (**Fig. S6C**). At shorter lag times, we observed a significant increase of approximately 100% in the PYR1->PV2 TE of the female KI network relative to its control for all peak Iapp values (**Fig. S6C**).

This increase in TE resulted in a significant reduction in both the mean (**Fig. S6D**) and standard deviation (**Fig. S6E**) of the PYR1xPV2 latency in the female KI network compared to controls, suggesting that the sculpting of the PV2 firing pattern is more precise and consistent in the female KI network than in the female WT network. The net effect of this result is a firing pattern of PV2 that is more effective in suppressing the excitation reaching PYR2 (**Fig. S6F**).

In the female KI network, low intensity I_app_ did not alter PYR1->PYR2 TE relative to controls, suggesting intact PYR model cell communication (**Fig. 7E**). However, at a higher I_app_, we observed significant changes in the PYR1->PYR2 TE (**Fig. 7E**, *right*), suggesting that neural communication in the female KI network is slightly altered with stronger stimulation compared to controls. These results also suggest that female PYR model cells are more resilient to inhibitory circuit perturbations than male networks.

In total, our results demonstrate that changes in intrinsic properties and synaptic connectivity associated with PV-mC4-OE decrease synaptic fidelity between model PYR and PV cells and cause hyperexcitability in a network model of male mice with increased levels of *mC4* in PV cells.

## DISCUSSION

Using a new model to conditionally overexpress *mC4*, we have discovered that mPFC PV cells in male mice are susceptible to complement dysfunction. Additionally, we have established a connection between the *mC4*-driven alterations in the circuitry of the mPFC and pathological anxiety-like behavior in male mice. Increased levels of *mC4* in PV neurons also disrupted both excitatory and inhibitory inputs to fast-spiking cells in male but not female mice. Furthermore, we have demonstrated that specific OE of *mC4* in PV cells led to opposing effects on the excitability of cortical cells. While mC4-OE in PV cells drove a decrease in the excitability of both male fast-spiking cells and PYRs, it led to hyperexcitability of female PV cells. By utilizing a Cre-driver line to induce mC4*-*OE in all neurons, we also observed that specific *mC4* dysfunction in PV cells has a greater adverse effect on anxiety-like behavior than widespread neuronal complement alterations. Using a simple computational model, we demonstrated that PV-mC4-driven inhibitory microcircuit deficits in the male model network led to disrupted neural communication between PYR model cells and hyperexcitability. Overall, these results establish a causative link between the SCZ-associated gene *C4* and the vulnerability of fast-spiking cells, which are crucial for the function of the mPFC.

### Synaptic alterations in fast-spiking cells with PV-specific *mC4* overexpression

Here, we demonstrate that in male mice, conditionally targeting mC4-OE to PV cells leads to a drastic loss of excitatory drive on this inhibitory cell type that is accompanied by increased inhibition. Several lines of evidence point to synaptic dysfunction and pathological excitatory synaptic loss as prominent features of SCZ (22,98–100). In support of this, our group and others previously demonstrated that increased levels of *C4* in developing L2/3 mPFC PYRs is sufficient to cause a significant loss of excitatory synapses, leading to mPFC circuit dysfunction (34,101). Our results reveal a significant decrease in the frequency of mEPSCs on PV cells without alterations in their amplitude. Although the underlying mechanism is not clear, our results suggest that the decline in excitatory drive to fast-spiking cells is either a reduction in the probability of presynaptic release or synapse number. In conjunction with previous results (34,101), this suggests that synapse loss is the most likely mechanism of hypoconnectivity in fast-spiking neurons.

While it is yet to be established whether pathological dysfunction in SCZ is confined to specific microcircuitry, a study conducted in SCZ post-mortem tissue demonstrated that there is a decrease of excitatory synapses on PFC PV cells relative to control subjects (22). In support of this, dysregulated ErbB4, a receptor of the SCZ-linked protein neuregulin-1, may contribute to lower activity of PV cells by reducing their excitatory inputs (102). A decrease in the excitatory drive to fast-spiking interneurons has also been observed in mouse models of AD (103,104) and neurodevelopmental disorders (105–107), suggesting that dysfunction in feed-forward excitatory synapses to fast-spiking cells is a common denominator in brain pathology.

A long-standing hypothesis is that defects in the GABAergic inhibitory system can contribute to SCZ (108). Additionally, cognitive impairment in SCZ could be the result of dysfunction in the convergence of glutamatergic and GABAergic systems (109). One possible outcome of decreased excitation on PV cells in the male PV-mC4-KI mouse is a disruption in the dynamics of excitation and inhibition, tipping the scales towards the side of unchecked excitation and excess glutamatergic release. This also aligns well with the NMDA-hypofunction SCZ model (110–112), where the loss of NMDA receptors, specifically on interneurons, results in hypoactivity of PV neurons. Alterations in inhibitory circuitry could also alter the timing of excitation and inhibition (113) that controls oscillatory activity and information flow (10).

We observed a significant increase in the amplitude of mIPSCs in PV cells in PV-mC4-KI male, but not female mice, suggesting an enhancement of inhibitory inputs to fast-spiking cells.

Naturally, this effect would amplify the putative decrease in PV cell activity in the male PV-mC4-KI mice, already caused by the reduction in the excitatory drive to this interneuron. As increased inhibition of PV cells is counterintuitive to the effects that increased complement activity would have on inhibitory synapses or a compensatory change to enhance the drive of fast-spiking cells, we can only conclude that these are *mC4*-driven maladaptive changes in the male brain. Whether this increase in mIPSC amplitude is driven by presynaptic changes in quantal size or postsynaptic changes in GABA receptor subunit composition or sensitivity is unknown and will require deeper investigation.

Microglia-dependent synaptic engulfment is an established mechanism for complement-driven synaptic loss in the normal and diseased brain (49,114–116). Studies using mice that lack specific complement genes have shown that these immune molecules contribute to synaptic plasticity (117–119). In fact, complement upregulation has been observed in several neurodegenerative diseases where synaptic loss is a prominent feature (120–122). A recent study also showed that C1q, the initiating member of the classic complement pathway, binds neuronal activity-regulated pentraxin (Nptx2) (123), an immediate early gene highly enriched at excitatory synapses on PV cells, where we observed the most drastic phenotype. Furthermore, deletion of Nptx2 caused increased activation of the classical complement pathway and microglia-mediated elimination of excitatory synapses on PV cells (123), supporting this established mechanism of synaptic loss in excitatory inputs on PV neurons. Still, other non-glia mechanisms could underlie excitatory synaptic loss in interneurons. In support of this, our group (124) used STED imaging in mPFC slices (125) to demonstrate that increased levels of *mC4* accelerate the accumulation of the postsynaptic receptor GluR1 in neuronal LAMP1-positive lysosomes, leading to pathological synaptic loss.

### The mPFC and neuropsychiatric disorders

The lifetime prevalence of anxiety disorders is close to 30% (126) and it is highly comorbid with other neuropsychiatric disorders, including SCZ (90,127,128). Our approach of conditionally targeting *mC4* in fast-spiking cells provides a unique example that establishes a causal relation between elevated levels of the SCZ-associated gene *C4* in these cells and enhanced anxiety-like behavior and mPFC circuit dysfunction in male mice, shedding light on the intricate dynamics of neuropsychiatric disorders.

We have focused on the mPFC to establish a connection between altered circuitry and disrupted emotional behavior. Genetic insults and chronic stress have lasting effects on the PFC that lead to alterations in cognitive and social function (129–132). In the mouse mPFC, inhibitory neurons respond to a variety of social and emotional stimuli (59,133). Additionally, PV cells coordinate and enhance the neuronal activity of PFC projection neurons to drive fear expression in the mouse (134). Consistent with its function in regulating emotional behavior, we observed that increased levels of *mC4* in PV neurons lead to synaptic alterations in fast-spiking cells and opposing effects in the excitability of cortical cells in male and female mice.

The PFC is plays a crucial role in social cognition, enabling us to understand and interpret the actions of others, which is fundamental for effective social interaction (135,136). Here we show that while increased levels of *mC4* in PV cells did not cause a drastic deficit in social behavior, overexpressing mice exhibited deficits in subclasses of exploratory social behavior, linking defects in inhibitory circuits to the initiation of social behaviors. In support of the role of mPFC PV neurons in the regulation of social behavior, Bicks et al. (137) demonstrated that PV cell activity in the mPFC preceded an active social episode, or an episode initiated by the experimental mouse. Similarly, we showed that increased levels of *mC4* in PV cells lead to deficits in active but not passive social interactions. Finally, in contrast to the deficits in anxiety-like behavior, *mC4*-driven social deficits were not sexually dimorphic, suggesting that in mice social deficits might have a distinct etiology from pathological anxiety-like behavior.

### Complement dysfunction and the mPFC

Although there is a strong link between immune dysfunction and neuropsychiatric disorders (49,138–143), more research is needed to establish a connection between complement dysfunction and specific circuitry underlying emotional behavior. Disruption of Csmd1, which is a C4 inhibitor, induces behaviors reminiscent of blunted emotional responses, anxiety, and depression (144). Additionally, Crider et al. (145) found a significant increase in C3 expression, a downstream effector of C4, in the PFC of depressed suicide subjects. Together with previous results (34,101), we showed that *mC4* alterations in specific cell-types are linked to mPFC-related pathologies. In summary, these results suggest that the PFC is a brain region susceptible to pathological complement activity.

### Increased levels of *mC4* in PV neurons, cortical function, and sex-related pathologies

Using a new mouse model, we show that targeted OE of *mC4* specifically in fast-spiking cells induces pathological anxiety-like behavior in male mice while sparing females. In the male mPFC, this sex-specific behavioral change correlates with a decrease in excitatory synaptic inputs to fast-spiking neurons, coupled with an increase in their inhibitory synapses, potentially resulting in reduced activity of this interneuron. During development, the maturation of fast-spiking cells contributes to the wiring of the neural networks, controlling the critical window of plasticity (146–149). Therefore, alterations in the developmental plasticity windows driven by increased levels of *mC4* in PV cells may cause the synaptic and excitability deficits we observed in the mPFC. There are also sex-dependent differences in the developmental cortical mechanisms of plasticity (150), which are regulated, in part, by PV cell activity, including their feed-forward circuits (147,148). Therefore, increased levels of *mC4* in fast-spiking cells could alter the function of the mPFC through distinct mechanisms in males and females, explaining the sex-divergent outcomes.

### Alterations in temporal fidelity and neural communication in SCZ and other neuropsychiatric disease

Using simulations, we showed a significant decrease in the firing rate of PV model cells and hyperactivity of PYR model cells in the male KI network compared to the male WT network. More critically, we also showed a significant broadening of the lag times over which activity in the presynaptic PYR influences the postsynaptic PYR, a product of defunct inhibition. This deficit in inhibition was evident in the drastic reduction of firing rates of PV2 in the male KI network and was bolstered by a reduction in the PYR1->PV2 transfer entropy compared to that of the male WT network.

During development, neuronal networks are far from static; an ever-dynamic landscape, synaptic connections are constantly being formed, lost, strengthened, or weakened across development (151–153). The fundamental unit of this plasticity is the information transferred from the presynaptic partner to its postsynaptic partner(s), the efficacy or existence of this connection being largely an activity-dependent factor (152,154). Moreover, disruptions in spike-timing and consequential deficits in synaptic plasticity are core features of neuropsychiatric diseases, including SCZ (29,83,155–157).

Thus, the specific finding of increased transfer entropy across a broader lag-time window in the male KI network is of particular importance because it suggests a major disruption in the temporal precision of effective information flow. In support of this, disruption of SCZ-associated risk genes that specifically contribute to spike-timing and plasticity in animal models have been shown to evoke a broad spectrum of SCZ-associated behavioral and synaptic deficits (158–161).

In the case of the female WT and KI networks, a seemingly-incompatible result emerged whereby, despite a lack of change in PV2 firing rate, the firing rate of PYR2 was reduced in the KI female condition. However, we observed a significant increase in the PYR1->PV2 transfer entropy in the female KI network that, despite not resulting in a change in the overall number of spikes evoked in PV2, shaped the temporal sequence of when those spikes of PV2 occurred more precisely. As a result, the effective inhibition experienced by PYR2 aligned more strongly with the excitation to PYR2 (from PYR1) in time, leading to an overall decrease in PYR2 firing rate.

Relevant to several broader hypotheses of SCZ is the convergence of altered neural communication and temporal fidelity. Using the specific experimental changes in intrinsic excitability and synaptic connectivity associated with PV-mC4-OE to guide our computational model, we too were led to exactly this same point of convergence. From the decades-enduring synaptic hypothesis of SCZ first proposed by Irwin Feinberg (98) to modern hypotheses of NMDAR hypofunction (111,156,162), many of the dominating views of SCZ pathogenesis feature disruptions in spike-timing and resultant deficits in the effective flow of information from one neuron to the next. Moreover, computational models of brain dysfunction relevant to SCZ from other groups consistently converge on this same feature (156).

Separately, Murray et al (163) implemented synaptic disinhibition – similar to our model of PV hypofunction – in a local-circuit PFC network model and found that this manipulation caused an increase in PYR firing rate (as in our model), broadened network activity, and decreased memory precision. Overall, our simulation provides evidence that a genetic alteration in fast-spiking cells leads to unique sex-dependent phenotypes in a model network, highlighting how cellular and synaptic phenotypes interact to produce complex neural network deficits in diseased states.

### Weaknesses of this study

We used a unique genetic approach to increase levels of *mC4* in PV cells globally. However, besides the PFC, we did not include recordings in other brain regions related to emotional regulation. Alterations in the inhibitory microcircuitry of other anxiety-related areas may underlie the behavioral effects that we have captured (164,165). Nevertheless, our findings demonstrate that conditional overexpression of *mC4* in fast-spiking cells results in synaptic and excitability deficits that are consistent with the role of mPFC PV cells in regulating emotional behavior.

SCZ is a complex disorder and it’s likely that multiple genetic and non-genetic factors contribute to its pathogenesis, each potentially impacting synaptic function and the excitability of cortical cells in different but converging ways. In light of this caveat, we provide a new mouse model where complement dysfunction in PV cells causes cellular and behavioral dysfunction reminiscent of PFC-associated neurological conditions.

While we used a computational model to better understand how the experimental results we gathered may cause network-level deficits in an intact system, our model has several limitations. First, the model has a simple architecture with strictly feed-forward connections. Thus, our ability to capture complex network interactions was limited. Moreover, the model was composed of only four interconnected model neuron units, a drastic simplification of the rich connectivity profiles of these cells in the intact rodent brain. Finally, the weights for all synaptic connections were altered in accordance with relative changes observed in the experimental mPSC data recorded in PYRs and PV cells in acute brain slices. However, these changes might not reflect *in vivo* properties. Nevertheless, despite these simplifications and limitations, our simulation of a network with increased levels of *mC4* in male fast-spiking cells is consistent with previous models of disrupted neuronal communication and prefrontal circuit dysfunction in schizophrenia (83).

### Concluding Remarks

Here, we have generated a unique mouse model to overexpress *C4* conditionally. Diseases linked to increased *C4* levels often have autoimmune or inflammatory aspects. Therefore, this mouse can be used to target specific cell types and tissues to determine the role of this important gene in various diseases outside of the nervous system or test the efficacy of pharmacology to target complement-related diseases. Together with previous studies, we have established *C4* as an important regulator of pathological synaptic loss in the prefrontal cortex, a region associated with several neuropsychiatric disorders. Furthermore, by conditionally overexpressing *C4* in fast-spiking cells, we have identified a connection between dysfunction of inhibitory circuits in the prefrontal cortex and pathological anxiety-like behavior in male mice.

## Supporting information

Table with statistics

## ACKNOWLEDGEMENTS

We would like to thank Dr. Todd Blute and the Boston University Biology Imaging Core for providing support for the confocal microscope. We thank members of the Cruz-Martín lab for critical reading of the manuscript and helpful discussions. This work was supported by a National Institutes of Health R01 (NIMH, 5R01MH129732-02) and an industry grant (to generate mC4-KI mouse, Biogen, #55206943) to A.C-M., a Brenton R. Lutz Award to R.A.P, a SURF/NSF-REU program (NSF, REU grant IOS-1659605) to J.R.G and N.M.P.L., and an NSF Award #2319321 to K.S.

## AUTHOR CONTRIBUTIONS

L.A.F. and A.C-M. conceptualized experiments including formulating composition, goals, and scope of the paper and approaches for analyses. L.A.F., R.A.P., M.S., A.B., S.B., J.R.G and N.M.P.L. collected the data and performed experiments. L.A.F. and R.A.P. performed data curation. L.A.F., R.A.P., A.B., M.S., and A.C.M analyzed data. L.A.F. and R.A.P. contributed code for data analysis. L.A.F. and A.C-M. contributed to parts of the original draft, including figure design and generation. L.A.F. developed the computational model with the support of J.C.N. and K.S. All authors contributed to revision and editing of the draft. A.C-M. obtained funding and supervised the project providing mentorship, oversight, and project administration.

## DECLARATION OF GENERATIVE AI AND AI-ASSISTED TECHNOLOGIES

The authors declare that no AI or AI-assisted technologies were used in the writing process.

## DATA AVAILABILITY

Data are available at https://osf.io/je38k/

## CODE AVAILABILITY

Custom-written routines are available at https://github.com/CruzMartinLab

## SUPPLEMENTAL FIGURE AND TABLE LEGENDS

**Supplemental Figure 1.**
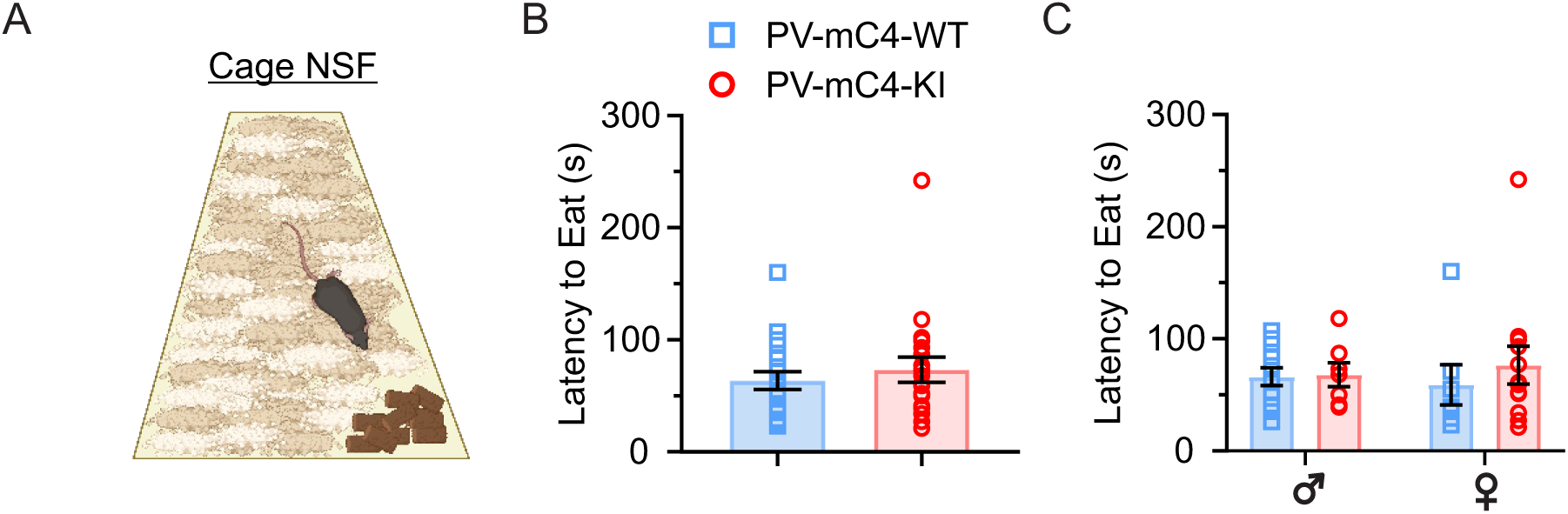
No change in latency to feed in the Cage NSF in PV-mC4-KI mice relative to controls. (**A**) Schematic of the Cage NSF. Latency to feed (s) was measured when WT or KI mice were placed in a more familiar environment, a standard cage. (**B**) No change in the latency to feed between WT and KI mice (Mann-Whitney test, *p*=0.5881). (**C**) No change in the latency to feed between groups, separated by sex (Two-way ANOVA, Condition x Sex: F(_1,34_)=0.2861, *p*=0.5962. Condition: F(_1,34_)=0.4357, *p*=0.5137. Sex: F(_1,34_)=0.00207, *p*=0.9640). WT: blue squares, *N*=12 males, *N*=7 females. KI: red circles, *N*=7 males, *N*=12 females. For all statistics,\**p*<0.05. Mean ± SEM shown.

**Supplemental Figure 2.**
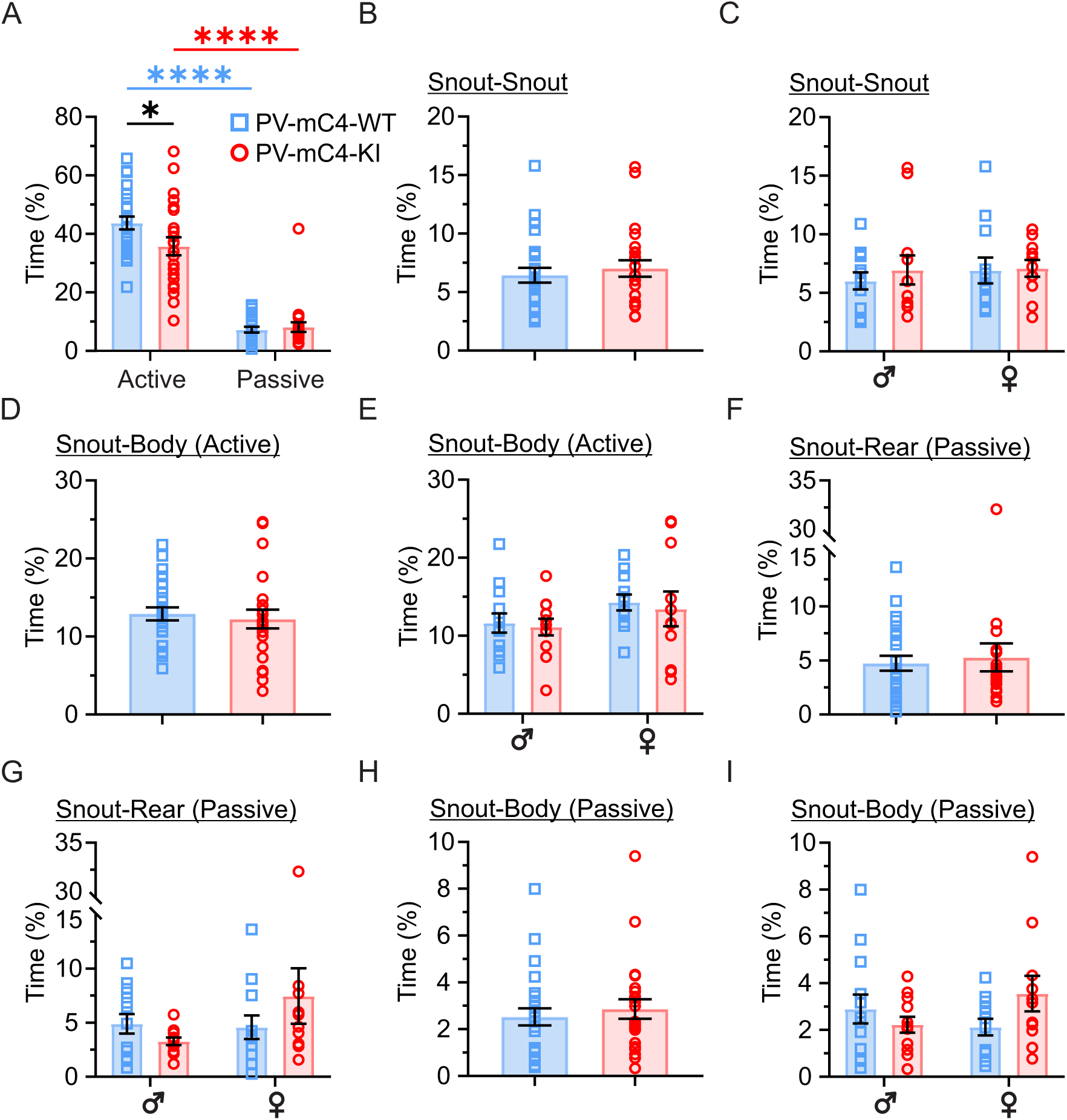
No changes in less-frequent sub-classes of social behavior with increased levels of *mC4* in PV cells. (**A**) Both WT and KI mice initiate most interactions during the juvenile interaction task (Condition x Interaction class: F(_1,92_)=4.484, \**p*=0.0369. Condition: F(_1,92_)=2.881, *p*=0.0930. Interaction class: F(_1,92_)=236.1, \*\*\*\**p*<0.0001. Post-test: Active vs. Passive WT, \*\*\*\**p*<0.0001, KI, \*\*\*\**p*<0.0001) with KI mice engaging less in active interactions compared to controls (WT vs. KI, \**p*=0.0488). (**B**, **C**) No change in reciprocal snout-snout interactions (13% of total social interactions) in KI mice, compared to controls (B: Mann-Whitney test, *p*=0.4993. C: Condition x Sex: F(_1,44_)=0.1482, *p*=0.7021. Condition: F(_1,44_)=0.3293, *p*=0.5690. Sex: F(_1,44_)=0.2833, *p*=0.5972). (**D**, **E**) No change in the active snout-body interaction (23% of total social interactions) in KI mice (D: t-test with Welch’s correction, *p*=0.6501. E: Condition x Sex: F(_1,44_)=0.0133, *p*=0.9088. Condition: F(_1,44_)=0.2210, *p*=0.6406. Sex: F(_1,44_)=3.028, *p*=0.0888). (**F**, **G**) No differences in the passive snout-rear interaction (9% of total social interactions) between groups (F: Mann-Whitney test, *p*=0.9186. G: Condition x Sex: F(_1,44_)=2.601, *p*=0.1139. Condition: F(_1,44_)=0.2085, *p*=0.6502. Sex: F(_1,44_)=1.930, *p*=0.1718). (**H**, **I**) No differences in the passive snout-body interaction (5% of total social interactions) between groups (H: Mann-Whitney test, *p*=0.5392, I: Condition x Sex: F(_1,44_)=3.745, *p*=0.0594. Condition: F(_1,44_)=0.4934, *p*=0.4861. Sex: F(_1,44_)=0.2645, *p*=0.6096). WT: blue squares, *N*=13 males, *N*=12 females. KI: red circles, *N*=12 males, *N*=11 females. For all statistics,\**p*<0.05, \*\**p*<0.01, \*\*\**p*<0.001, \*\*\*\**p*<0.0001. Two-way ANOVA, unless otherwise stated. Mean ± SEM shown.

**Supplemental Figure 3.**
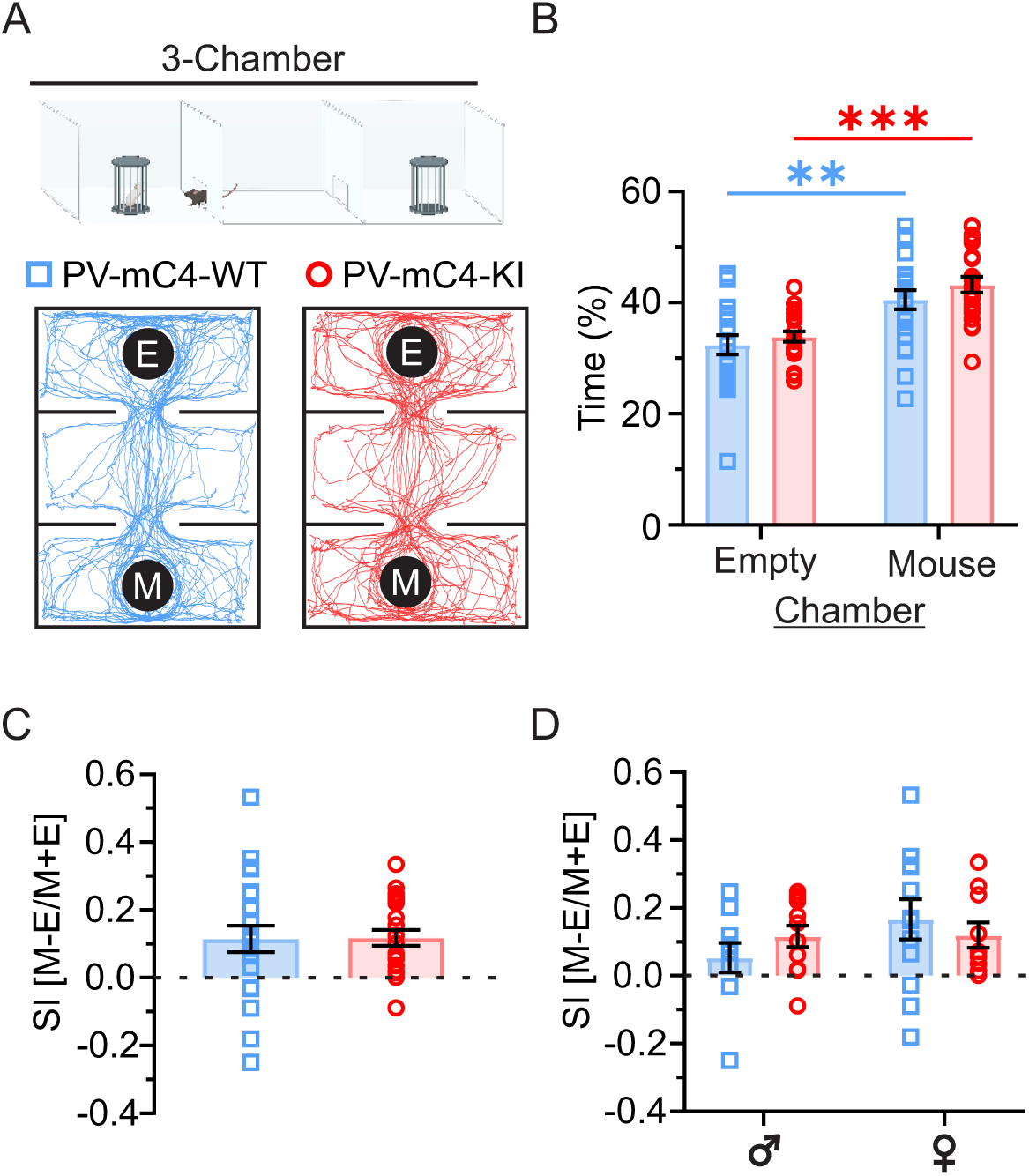
Overexpression of *mC4* in PV cells did not alter social interactions in the three-chamber assay. (**A**) (*Top*) Schematic representation of three-chamber assay. (*Bottom*) Representative trajectories (tracked with DLC) of WT (*blue*) and KI *(red*) mice in the three-chamber assay. E, *empty cup*; M, *mouse cup*. (**B**) WT and KI mice spent more time in the chamber containing the mouse cup compared to the empty-cup chamber, suggesting that they prefer social interactions (Condition x Chamber: F(_1,84_)=0.1733, *p*=0.6782. Condition: F(_1,84_)=1.911, *p*=0.1706. Chamber: F(_1,84_)=34.19, \*\*\*\**p*<0.0001. Post-test: Mouse vs. Empty, WT \*\**p*=0.0013, KI, \*\*\**p*=0.0002). (**C**, **D**) No change in the social discrimination index (SI) between WT and KI mice (C: t-test with Welch’s correction, *p*=0.9463. D: Condition x Sex: F(_1,40_)=1.455, *p*=0.2349. Condition: F(_1,40_)=0.03420, *p*=0.8542. Sex: F(_1,40_)=1.643, *p*=0.2074). WT: blue squares, *N*=10 males, *N*=12 females. KI: red circles, *N*=12 males, *N*=10 females. For all statistics,\**p*<0.05, \*\**p*<0.01, \*\*\**p*<0.001, Two-way ANOVA, unless otherwise stated. Mean ± SEM shown.

**Supplemental Figure 4.**
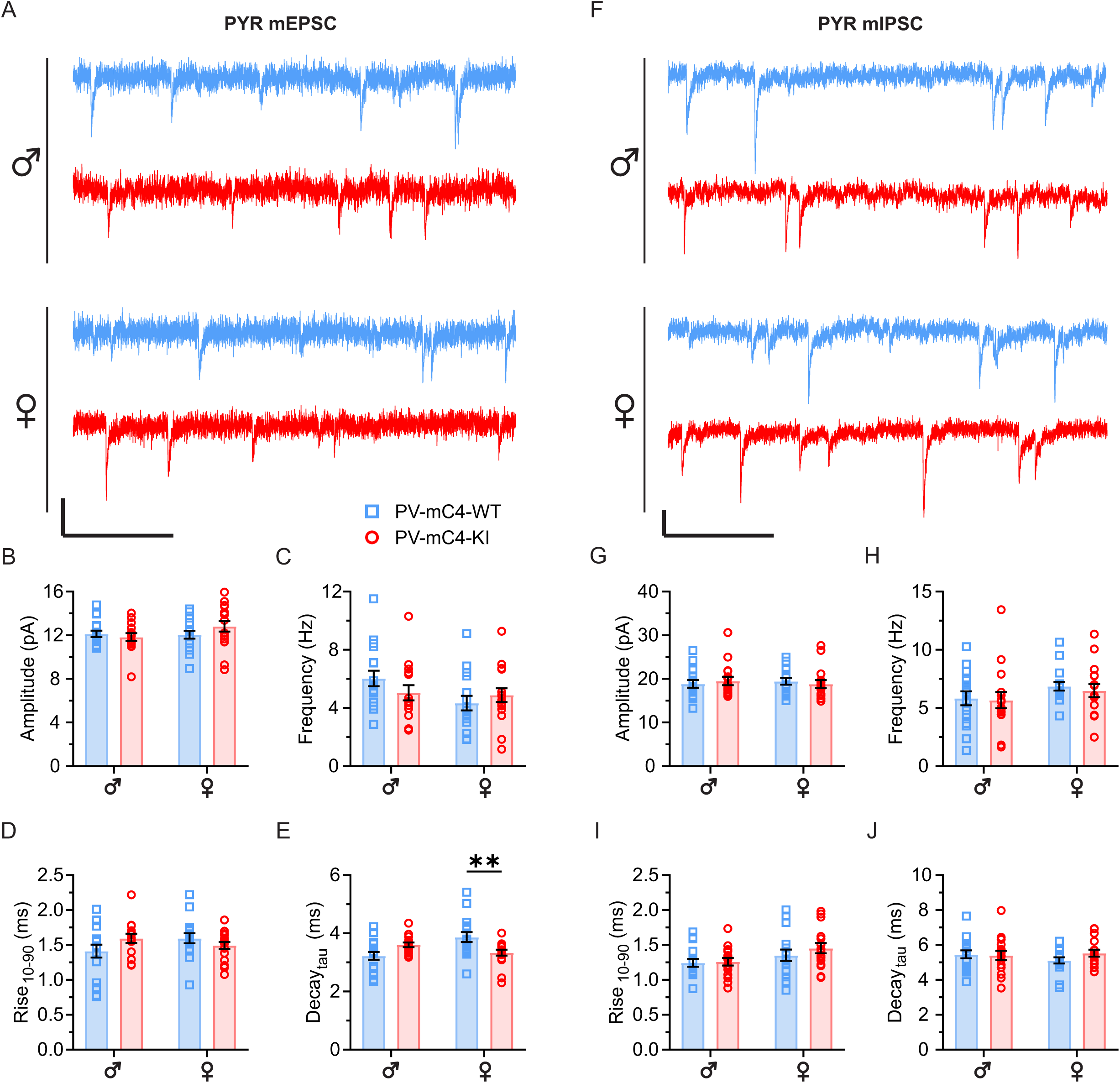
PV-specific mC4-OE alters the kinetics of mEPSCs in PYRs of female mice. (**A**) Representative 1 s traces of mEPSCs recorded in PYRs in P40-60 young adult WT (*blue*) and KI (*red*) mice, *scale bar* = 250 ms, 10 pA. (**B**) No change in mEPSC amplitude in KI mice, relative to controls (Condition x Sex: F(_1,61_)=1.945, *p*=0.1681. Condition: F(_1,61_)=0.4158, *p*=0.5214. Sex: F(_1,61_)=1.397, *p*=0.2418). (**C**) No change in in mEPSC frequency in KI mice (Condition x Sex: F(_1,61_)=2.291, *p*=0.1353. Condition: F(_1,61_)=0.1903, *p*=0.6642. Sex: F(_1,61_)=3.341, *p*=0.0725). (**D**) mC4-OE did not alter mEPSC rise Rise_10-90_ (Condition x Sex: F(_1,61_)=3.916, *p*=0.0524. Condition: F(_1,61_)=0.3104, *p*=0.5795. Sex: F(_1,61_)=0.3120, *p*=0.5785). (**E**) Increased expression of *mC4* led to a significant decrease in mEPSC Decay_tau_ in KI females (Condition x Sex: F(_1,61_)=12.74, \*\*\**p*=0.0007. Condition: F(_1,61_)=0.3684, *p*=0.5461. Sex: F(_1,61_)=2.183, *p*=0.1447. Post-test: WT vs. KI males, *p*=0.0822, females, \*\**p*=0.0083). (**F**) Representative 1 s traces of mIPSCs recorded in PYRs in young adult WT (*blue*) and KI (*red*) mice, *scale bar* = 250 ms, 10 pA. (**G**) No change in mIPSC amplitude in KI mice, relative to controls (Condition x Sex: F(_1,60_)=0.5354, *p*=0.4672. Condition: F(_1,60_)=0.00013, *p*=0.9909. Sex: F(_1,60_)=0.00367, *p*=0.9519). (**H**) mIPSC frequency was not different between groups (Condition x Sex: F(_1,60_)=0.03569, *p*=0.8508. Condition: F(_1,60_)=0.2141, *p*=0.6452. Sex: F(_1,60_)=2.690, *p*=0.1062). (**I**) Increased expression of *mC4* did not impact mIPSC Rise_10-90_ (Condition x Sex: F(_1,60_)=0.3837, *p*=0.5380. Condition: F(_1,60_)=0.7667, *p*=0.3847. Sex: F(_1,60_)=5.047, \**p*=0.0284. Post-test: WT vs. KI males, *p*=0.9795, females, *p*=0.5025). (**J**) Decay_tau_ was not changed in KI mice (Condition x Sex: F(_1,60_)=1.153, *p*=0.2871. Condition: F(_1,60_)=0.7059, *p*=0.4042. Sex: F(_1,60_)=0.2624, *p*=0.6104). WT: blue squares. KI: red circles. *N* represents cells. mEPSC WT *N*=17 males, *N*=16 females; mEPSC KI: *N*=15 males, *N*=17 females. mIPSC WT *N*=16 males, *N*=16 females; mIPSC KI: *N*=16 males, *N*=16 females. For all statistics,\**p*<0.05, \*\**p* <0.01, Two-way ANOVA, unless otherwise stated. Mean ± SEM shown.

**Supplemental Figure 5.**
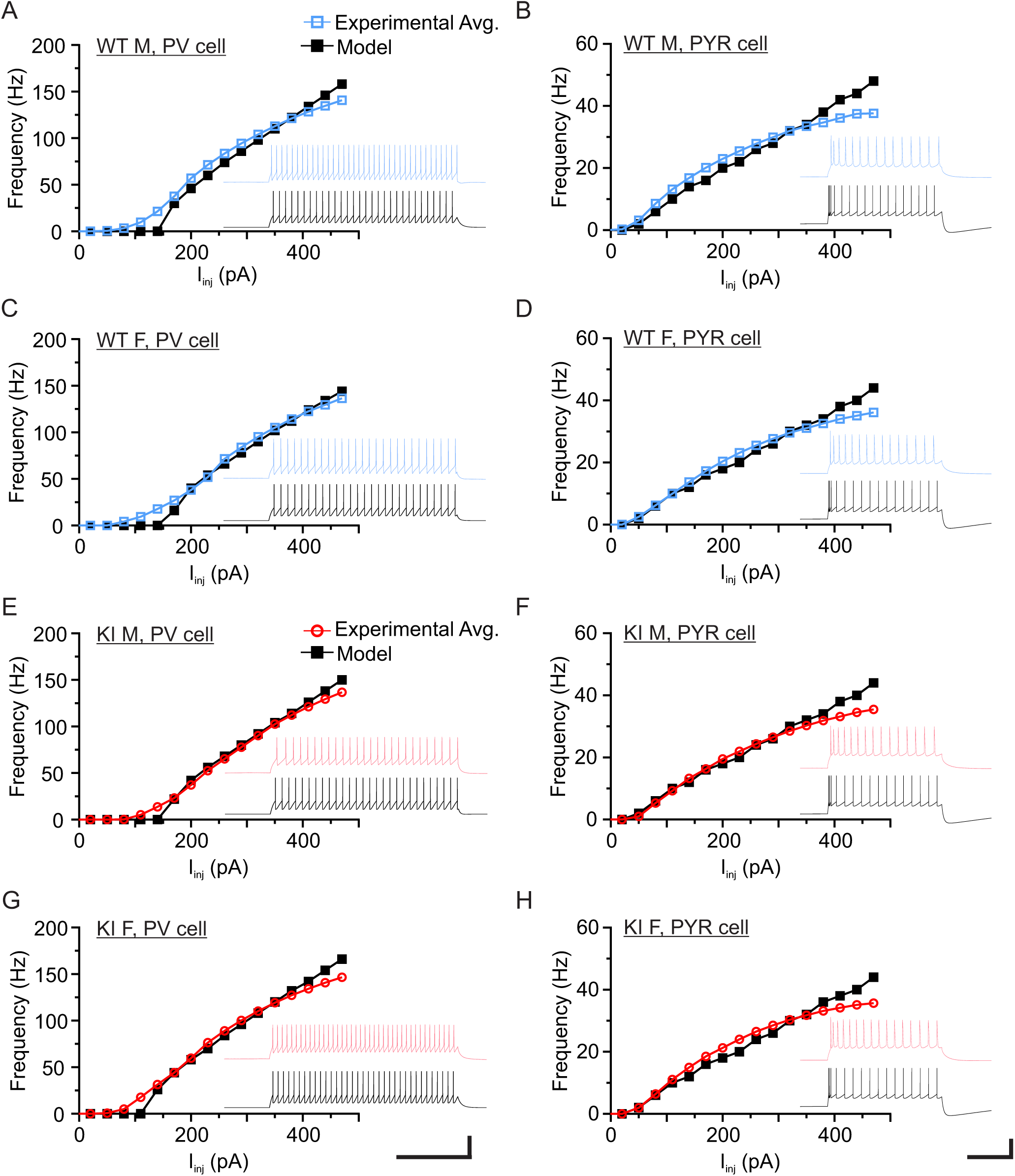
Modeled PV and PYR units have similar firing rates as experimental PV and PYR cells in acute brain slices. For all frequency vs. current (FI curve) plots, average experimental FI curve data (colored curve) is plotted against the equivalent FI curve for the respective modeled cell (black curve). All insets show a representative voltage trace from experimental data (colored voltage trace, top) and the voltage trace of the equivalent modeled cell (black voltage trace, bottom). Inset traces for experimental PV cells and modeled PV units (**A, C, E,** and **G**) are in response to an identical 230 pA square pulse (500 ms). Inset traces for experimental PYR and modeled PYR units (**B, D, F,** and **H**) are in response to an identical 350 pA square current pulse (500 ms). Scale for all insets, 200 ms/50 mV. (**A**, **B**) FI curve comparisons for WT male PV cells (**A**) and PYR (**B**). (**C**, **D**) FI curve comparisons for WT female PV cells (**C**) and PYR (**D**). (**E**, **F**) FI curve comparisons for KI male PV cells (**E**) and PYR (**F**). (**G**, **H**) FI curve comparisons for KI female PV cells (**G**) and PYR (**H**).

**Supplemental Figure 6.**
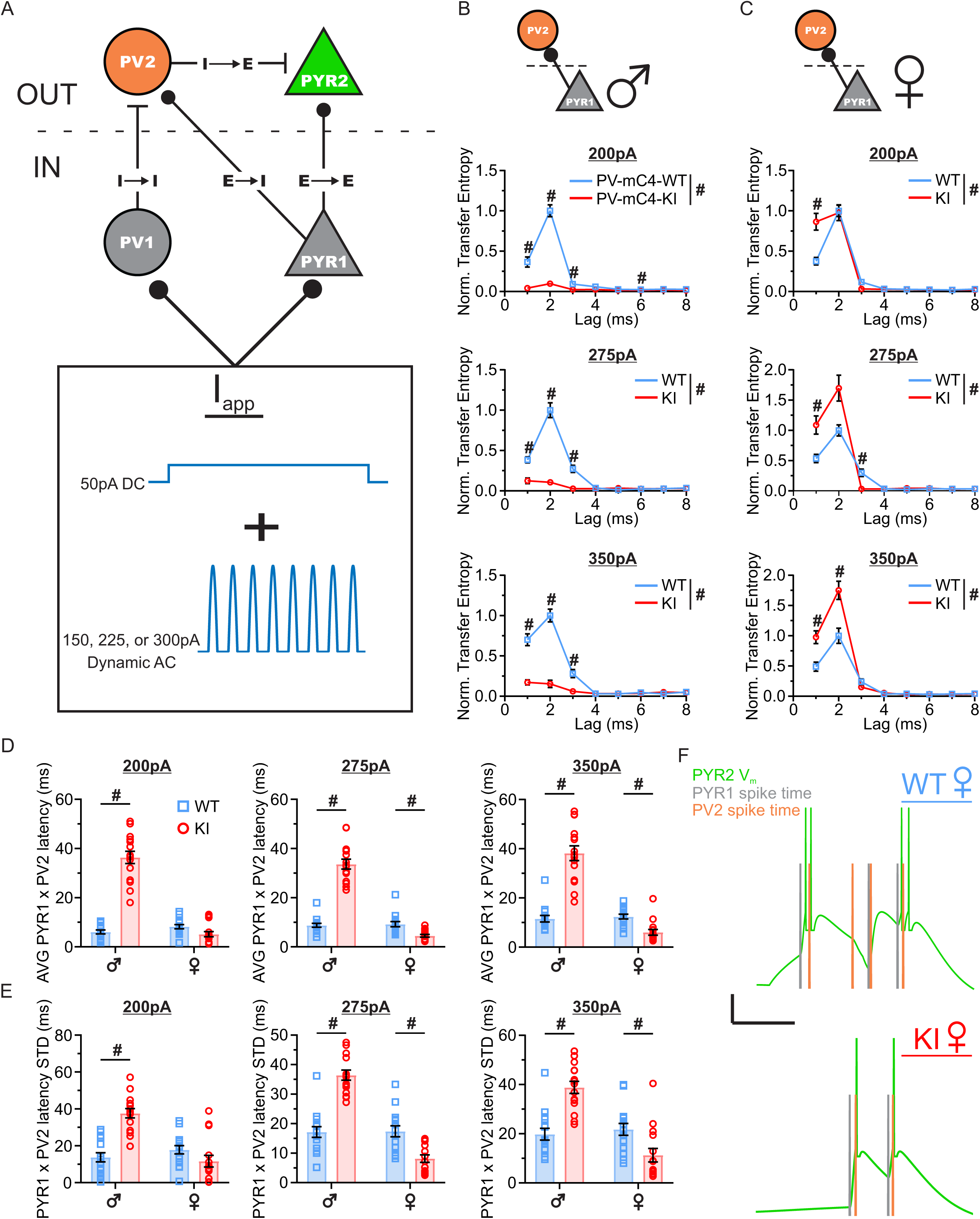
A computational model reveals that changes associated with PV-mC4-OE drive changes in PYR-PV information flow. (**A**) Schematic showing the network architecture and applied current (*Iapp*) of the computational model. (**B**) Plots showing the average transfer entropy (TE) at each delay (lag) of PYR1 onto PV2 in male WT (*blue*) and male KI (*red*) networks at peak *I_app_* of 200 pA (*top,* Condition x Lag: F(_7,196_)=67.00, \*\*\*\**p*<0.0001. Condition: F(_1,28_)=448.8, \*\*\*\**p*<0.0001. Lag: F(_1.541,43.16_)=93.00, \*\*\*\**p*<0.0001), 275 pA (*middle,* Condition x Lag: F(_7,196_)=51.27, \*\*\*\**p*<0.0001. Condition: F(_1,28_)=339.8, \*\*\*\**p*<0.0001. Lag: F(_1.603,44.90_)=73.85, \*\*\*\**p*<0.0001), and 350 pA (*bottom,* Condition x Lag: F(_7,196_)=42.06, \*\*\*\**p*<0.0001. Condition: F(_1,28_)=307.0, \*\*\*\**p*<0.0001. Lag: F(_2.123,59.44_)=73.98, \*\*\*\**p*<0.0001). (**C**) Plots showing the average transfer entropy (TE) at each delay (lag) of PYR1 onto PV2 in female WT (*blue*) and female KI (*red*) networks at peak *I_app_* of 200pA (*top*, Condition x Lag: F(_7,196_)=8.426, \*\*\*\**p*<0.0001. Condition: F(_1,28_)=10.37, \*\**p*=0.0032. Lag: F(_1.507, 42.19_)=141.1, \*\*\*\**p*<0.0001), 275 pA (*middle*, Condition x Lag: F(_7,196_)=8.855, \*\*\*\**p*<0.0001. Condition: F(_1,28_)=41.96, \*\*\*\**p*<0.0001. Lag: F(_1.367,38.27_)=83.36, \*\*\*\**p*<0.0001), and 350 pA (*bottom*, Condition x Lag: F(_7,196_)= 11.66, \*\*\*\**p*<0.0001. Condition: F(_1,28_)=43.53, \*\*\*\**p*<0.0001. Lag: F(_1.567,43.87_)=122.8, \*\*\*\**p*<0.0001). (**D**) The average latency from one PYR1 spike to the next soonest PV2 spike in WT and KI networks at peak *I_app_* of 200 pA (*left*, Condition x Sex: F(_1,56_)=127.8, \*\*\*\**p*<0.0001. Condition: F(_1,56_)=85.53, \*\*\*\**p*<0.0001. Sex: F(_1,56_)=96.97, \*\*\*\**p*<0.0001. Post-test: WT vs. KI males, \*\*\*\**p*<0.0001, females, *p*=0.2803), 275 pA (*middle*, Condition x Sex: F(_1,56_)=141.7, \*\*\*\**p*<0.0001. Condition: F(_1,56_)=65.75, \*\*\*\**p*<0.0001. Sex: F(_1,56_)=133.0, \*\*\*\**p*<0.0001. Post-test: WT vs. KI males, \*\*\*\**p*<0.0001, females, \**p*=0.0191), and 350 pA (*right*, Condition x Sex: F(_1,56_)=83.60, \*\*\*\**p*<0.0001. Condition: F(_1,56_)=31.45, \*\*\*\**p*<0.0001. Sex: F(_1,56_)=75.19, \*\*\*\**p*<0.0001. Post-test: WT vs. KI males, \*\*\*\**p*<0.0001, females, \**p*=0.0305). (**E**) The standard deviation of the latency from one PYR1 spike to the next soonest PV2 spike in WT and KI networks at peak *I_app_* of 200 pA (*left*, Condition x Sex: F(_1,56_)=32.54, \*\*\*\**p*<0.0001. Condition: F(_1,56_)=11.29, **p=0.0014. Sex: F(_1,56_)=17.23, \*\*\**p*=0.0001. Post-test: WT vs. KI males, \*\*\*\**p*<0.0001, females, *p*=0.1953), 275 pA (*middle*, Condition x Sex: F(_1,56_)=71.33, \*\*\*\**p*<0.0001. Condition: F(_1,56_)=8.837, \*\**p*=0.0043. Sex: F(_1,56_)=68.89, \*\*\*\**p*<0.0001. Post-test: WT vs. KI males, \*\*\*\**p*<0.0001, females, \*\*\**p*=0.0006), and 350 pA (*right*, Condition x Sex: F(_1,56_)=34.73, \*\*\*\**p*<0.0001. Condition: F(_1,56_)=2.936, *p*=0.0921. Sex: F(_1,56_)=26.08, \*\*\*\**p*<0.0001. Post-test: WT vs. KI males, \*\*\*\**p*<0.0001, females, \*\**p*=0.0091).WT network: blue squares. KI network: red circles. *N* represents simulations. *N*=15 for all networks. (**F**) Representative simulated traces of PYR2 membrane voltage (green) overlaid with the timing of spikes of PYR1 (gray) and PV2 (orange) in the female WT (*top*) and female KI (*bottom*) network, *scale bar* = 20 ms, 20mV. For all statistics, \**p*<0.05, \*\**p*<0.01, \*\*\**p*<0.001, \*\*\*\**p*<0.0001, Repeated-measure Two-way ANOVA with multiple comparisons (**B, C**) or Two-way ANOVA (**E, F**). Mean ± SEM shown. For all plots, ‘*#*’ indicates all *p*<0.05.

**Supplemental Table 1. Active and passive electrophysiological properties of PV cells in PV-mC4-WT and KI mice – Main Effects.** Table displays the main effects results of each Two-way ANOVA for PV cells. Significant main effects (*p*<0.05) are bolded.

**Supplemental Table 2. Active and passive electrophysiological properties of PV cells in PV-mC4-WT and KI mice – Post-tests.** Table displays WT and KI means ± SEM, and the associated *p*-value (Šídák’s multiple comparisons) for the comparison between conditions if applicable. Data are separated by sex, male data on the left and female data on the right. For all statistics \**p*<0.05, Two-way ANOVA. In all KI mice, resting V_m_ was significantly more depolarized in response to PV-mC4-OE, relative to WT (WT: -67.25 ± 0.8696 mV vs. KI: -64.05 ± 1.209 mV, t-test with Welch’s correction, \**p*=0.0356), data not shown in table.

**Supplemental Table 3. Active and passive electrophysiological properties of PYRs in PV-mC4-WT and KI mice – Main Effects.** Table displays the main effects results of each Two-way ANOVA for PYRs. Significant main effects (*p*<0.05) are bolded.

**Supplemental Table 4. Active and passive electrophysiological properties of PYRs in PV-mC4-WT and KI mice – Post-tests.** Table displays WT and KI means ± SEM, and the associated *p*-value (Šídák’s multiple comparisons) for the comparison between conditions if applicable. Data are separated by sex, male data on the left and female data on the right. For all statistics \**p*<0.05, Two-way ANOVA.

**Supplemental Table 5. PYR and PV cell parameters used in computational model.** DynaSim parameters. Specific values for E_L_, R_m_, C_m_, V_thresh_, and V_reset_ were determined experimentally.

**Supplemental Table 6. Synaptic parameters used in computational model.** DynaSim parameters. Synaptic connectivity conductance (g_syn_) was set to a default value of 0.03 for all connections, and was only altered where mEPSC or mIPSC frequency or amplitude recorded in PYR or PV cells was significantly different between groups across or within sex. Rise and decay kinetics for all parameters were determined experimentally.

