## Supplemental tables for "Overexpression of the schizophrenia risk gene C4 in PV cells drives sex-dependent behavioral deficits and circuit dysfunction"

| PV cell | Two-way ANOVA. $F_{(1,69)}$=, *p*= | | |
|---|---|---|---|
| | Sex x Condition | Sex | Condition |
| Resting $V_m$ (mV) | 0.0975, 0.7558 | 0.7875, 0.3779 | **4.309, *0.0416** |
| Input Resistance, $R_{in}$ (MΩ) | 2.925, 0.0917 | 2.057, 0.1560 | 0.2246, 0.6370 |
| Membrane Capacitance, $C_m$ (pF) | 0.5158, 0.4750 | 0.0619, 0.8042 | 2.926, 0.0917 |
| Decay constant, tau (ms) | 2.100, 0.1518 | **4.507, *0.0373** | 1.615, 0.2080 |
| $ISI_1/ISI_9$ | 0.0082, 0.9281 | 2.703, 0.1084 | 0.0327, 0.8570 |
| $ISI_4/ISI_9$ | 0.0826, 0.7748 | 0.9215, 0.3405 | 0.2781, 0.5997 |
| Sag Ratio, $F_{(1,60)}$ | 1.064, 0.3064 | 1.686, 0.1991 | 1.161, 0.2857 |

**Supplemental Table 1. Active and passive electrophysiological properties of PV cells in PV-mC4-WT and KI mice – Main Effects**

| PV cell | Male | | | Female | | |
|---|---|---|---|---|---|---|
| | Mean ± SEM | | post-test, p-value | Mean ± SEM | | post-test, p-value |
| | WT | KI | | WT | KI | |
| Resting $V_m$ (mV) | -68.11 ± 5.37 | -64.5 ± 7.23 | 0.1781 | -66.29 ± 5.02 | -63.63 ± 7.65 | 0.3916 |
| Input Resistance, $R_{in}$ (MΩ) | 161.27 ± 11.18 | 147.15 ± 9.36 | - | 158.11 ± 12.41 | 183.07 ± 12.38 | - |
| Membrane Capacitance, $C_m$ (pF) | 63.74 ± 2.58 | 67.52 ± 1.85 | - | 61.96 ± 3.64 | 71.20 ± 5.75 | - |
| Decay constant, tau (ms) | 9.78 ± 0.52 | 9.64 ± 0.58 | 0.9899 | 10.31 ± 0.93 | 12.44 ± 1.00 | 0.1171 |
| $ISI_1/ISI_9$ | 1.44 ± 0.42 | 1.52 ± 0.30 | - | 1.03 ± 0.01 | 1.05 ± 0.04 | - |
| $ISI_4/ISI_9$ | 0.98 ± 0.01 | 0.98 ± 0.02 | - | 1.00 ± 0.01 | 0.99 ± 0.01 | - |
| Sag Ratio | 0.040 ± 0.003 | 0.040 ± 0.004 | - | 0.049 ± 0.004 | 0.041 ± 0.004 | - |

**Supplemental Table 2. Active and passive electrophysiological properties of PV cells in PV-mC4-WT and KI mice – Post-tests.**

| PYR | Two-way ANOVA. $F_{(1,63)}$=, *p*= | | |
|---|---|---|---|
| | Sex x Condition | Sex | Condition |
| Resting $V_m$ (mV) | 0.0104, 0.9191 | 0.1306, 0.7190 | .0310, 0.8608 |
| Input Resistance, $R_{in}$ (MΩ) | 3.661, 0.0602 | 0.2326, 0.6313 | 0.1358, 0.7138 |
| Membrane Capacitance, $C_m$ (pF) | 0.1365, 0.7130 | **4.155, *0.0457** | 1.705, 0.1964 |
| Decay constant, tau (ms) | 2.826, 0.0977 | 0.6693, 0.4164 | 0.0719, 0.7894 |
| $ISI_1/ISI_9$ | 0.4917, 0.4857 | 1.259, 0.2660 | 0.3778, 0.5410 |
| $ISI_4/ISI_9$ | 1.396, 0.2419 | **4.856, *0.0312** | 0.0810, 0.7769 |
| Sag Ratio, $F_{(1,59)}$ | 2.894, 0.0942 | 0.1244, 0.7256 | 0.3072, 0.5815 |

**Supplemental Table 3. Active and passive electrophysiological properties of PYRs in PV-mC4-WT and KI mice – Main Effects.**

| PYR | Male | | | Female | | |
|---|---|---|---|---|---|---|
| | Mean ± SEM | | post-test, p-value | Mean ± SEM | | post-test, p-value |
| | WT | KI | | WT | KI | |
| Resting $V_m$ (mV) | -69.87 ± 1.75 | -70.0 ± 2.0 | - | -70.33 ± 1.66 | -70.83 ± 1.75 | - |
| Input Resistance, $R_{in}$ (MΩ) | 244.88 ± 15.80 | 204.61 ± 13.58 | - | 219.62 ± 19.20 | 246.89 ± 19.56 | - |
| Membrane Capacitance, $C_m$ (pF) | 145.91 ± 7.88 | 159.09 ± 11.02 | 0.4488 | 132.77 ± 5.35 | 140.14 ± 6.76 | 0.7437 |
| Decay constant, tau (ms) | 34.32 ± 1.92 | 31.22 ± 1.99 | - | 28.84 ± 2.50 | 33.16 ± 2.15 | - |
| $ISI_1/ISI_9$ | 0.32 ± 0.04 | 0.33 ± 0.02 | - | 0.37 ± 0.03 | 0.34 ± 0.03 | - |
| $ISI_4/ISI_9$ | 0.78 ± 0.02 | 0.81 ± 0.02 | 0.5393 | 0.84 ± 0.01 | 0.83 ± 0.02 | 0.7618 |
| Sag Ratio | 0.065 ± 0.005 | 0.075 ± 0.011 | - | 0.083 ± 0.009 | 0.063 ± 0.008 | - |

**Supplemental Table 4. Active and passive electrophysiological properties of PYRs in PV-mC4-WT and KI mice – Post-tests.**

| | PYR | | | | PV cell | | | |
|---|---|---|---|---|---|---|---|---|
| | Male | | Female | | Male | | Female | |
| | WT | KI | WT | KI | WT | KI | WT | KI |
| $E_L$ (mV) | -70.3 → | → | → | → | -67.2 | -64.1 | -67.2 | -64.1 |
| $R_m$ (MΩ) | 229.0 → | → | → | → | 162.4 → | → | → | → |
| $C_m$ (pF) | 144.5 → | → | → | → | 66.1 → | → | → | → |
| $V_{thresh}$ (mV) | -31.4 → | → | → | → | -37.0 | -34.0 | -34.9 → | → |
| $V_{reset}$ (mV) | -43.08 → | → | -44.7 | -41.5 | -54.0 → | → | → | → |
| $\tau_{sra}$ (ms) | 100 → | → | → | → | 5 → | → | → | → |
| $\Delta g_{sra}$ (nS/spk) | 0.00475 | 0.00525 → | → | → | 0.005 → | → | → | 0.007 |
| Integration Time Step (ms) | 0.1 → | → | → | → | → | → | → | → |

**Supplemental Table 5. PYR and PV cell parameters used in computational model.**

| | | $g_{syn}$ (nS) | $E_{syn}$ (mV) | Rise (ms) | Decay (ms) |
|---|---|---|---|---|---|
| PYR→PYR (E→E) | WT M | 0.03 | 0 | 1.52 | 3.41 |
| | KI M | | | | ↓ |
| | WT F | | | | 3.87 |
| | KI F | | | ↓ | 3.34 |
| PYR→PV (E→I) | WT M | ↓ | | 0.56 | 1.63 |
| | KI M | 0.0183 | | | |
| | WT F | 0.03 | | | |
| | KI F | | | ↓ | ↓ |
| PV→PYR (I→E) | WT M | | -81 | 1.33 | 5.37 |
| | KI M | | | | |
| | WT F | | | | |
| | KI F | | | ↓ | ↓ |
| PV→PV (I→I) | WT M | ↓ | | 0.83 | 3.44 |
| | KI M | 0.0348 | | | |
| | WT F | 0.03 | | | |
| | KI F | ↓ | ↓ | ↓ | ↓ |

**Supplemental Table 6. Synaptic parameters used in computational model.**
