## Supplementary material for "Overexpression of the schizophrenia risk gene C4 in PV cells drives sex-dependent behavioral deficits and circuit dysfunction": Table with statistics

| Figure 1. A novel transgenic mouse line permits PV cell specific overexpression of complement component 4. | | | | | |
|---|---|---|---|---|---|
| **Panel** | | | | | |
| **A** | N/A | | | | |
| **B** | N/A | | | | |
| **C** | **In both sexes, increased levels of mC4 in PV cells did not alter mouse weights compared to controls.** | | | | |
| | **Factor or comparison** | **Statistical test** | **F statistic** | **P value** | **Ns** |
| | Condition x Sex | Two-way ANOVA | F(1,44)=3.204 | *p*=0.0803 | |
| | Condition | Two-way ANOVA | F(1,44)=0.6311 | *p*=0.4312 | |
| | Sex | Two-way ANOVA | F(1,44)=103.2 | *********p*<0.0001** | |
| | **Post-test** | | | | |
| | WT vs. KI males | Šídák's multiple comparisons | N/A | *p*=0.9793 | WT, N=13, KI, N=12 |
| | WT vs. KI Females | Šídák's multiple comparisons | N/A | *p*=0.3947 | WT, N=12, KI, N=11 |
| **D** | **OE of mC4 in PV cells did not impact the distance traveled by mice in an open field** | | | | |
| | **Factor or comparison** | **Statistical test** | **F statistic** | **P value** | **Ns** |
| | Condition x Sex | Two-way ANOVA | F(1,34)=0.04903 | *p*=0.8261 | |
| | Condition | Two-way ANOVA | F(1,34)=0.004672 | *p*=0.9459 | |
| | Sex | Two-way ANOVA | F(1,34)=2.657 | *p*=0.1123 | |
| | | | | | WT, N=12, KI, N=7 |
| | | | | | WT, N=7, KI, N=12) |
| **E** | N/A | | | | |
| **F** | N/A | | | | |
| **G** | **No difference in PV cell density in the mPFC between WT and KI** | | | | |
| | **Factor or comparison** | **Statistical test** | **F statistic** | **P value** | **Ns** |
| | WT vs. KI | Mann-Whitney test | N/A | *p*=0.2319 | WT, N=8, KI N=7 |
| **H** | N/A | | | | |

| | | | | | |
|---|---|---|---|---|---|
| **I** | **In P21 KI mice, mC4 was significantly overexpressed in PV cells, but not in SST cells or all other DAPI-labeled cells** | | | | |
| | **Factor or comparison** | **Statistical test** | **F statistic** | **P value** | **Ns** |
| | Condition x Cell-type | Two-way ANOVA | F(2,18)=41.62, | ****p<0.0001 | |
| | Condition | Two-way ANOVA | F(1,18)=113.7 | ****p<0.0001 | |
| | Cell-type | Two-way ANOVA | F(2,18)=38.22 | ****p<0.0001 | |
| | **Post-test** | | | | |
| | WT vs. KI, PV | Šídák's multiple comparisons | | ****p<0.0001 | |
| | WT vs. KI, SST | Šídák's multiple comparisons | | *p*=0.1793 | |
| | WT vs. KI, Other | Šídák's multiple comparisons | | *p*=0.2166 | |
| | **In P21 KI mice, *mC4* expression was significantly greater in PV cells than in SST cells** | | | | |
| | PV vs. SST | Šídák's multiple comparisons | | ****p<0.0001 | |
| | PV vs. Other | Šídák's multiple comparisons | | ****p<0.0001 | |
| | | | | | WT *N*=5 at P21, KI N=3 at P21 |
| **J** | **In P65 mice, *mC4* was overexpressed in PV cells, but not in SST cells or all other DAPI-labeled cells** | | | | |
| | **Factor or comparison** | **Statistical test** | **F statistic** | **P value** | **Ns** |
| | Condition x Cell-type | Two-way ANOVA | F(2,24)=33.42 | ****p<0.0001 | |
| | Condition | Two-way ANOVA | F(1,24)=66.63 | ****p<0.0001 | |
| | Cell-type | Two-way ANOVA | F(2,24)=32.59 | ****p<0.0001 | |
| | **Post-test** | | | | |
| | WT vs. KI PV | Šídák's multiple comparisons | | ****p<0.0001 | |
| | WT vs. KI SST | Šídák's multiple comparisons | | p=0.6948 | |
| | WT vs. KI Other | Šídák's multiple comparisons | | p=0.7839 | |
| | **In KI mice, mC4 expression was significantly greater in PV cells than in SST cells** | | | | |
| | PV vs. SST | Šídák's multiple comparisons | | ****p<0.0001 | WT *N*=5 at P21, KI N=3 at P21 |
| | PV vs.Other | Šídák's multiple comparisons | | ****p<0.0001 | WT N=5 at P65, KI N=5 at P65 |

| | | | | | | | | | |
|---|---|---|---|---|---|---|---|---|---|
| **I, J** | **In PV-mC4-KI mice, the number of *mC4* puncta in PV cells was not different between P21 and 65** | | | | | | | | |
| | **Factor or comparison** | | **Statistical test** | | **F statistic** | | **P value** | | **Ns** |
| | Age x Cell-type | | Two-way ANOVA | | $F_{(2,18)}=2.454$ | | *p*=0.1142 | | |
| | Age | | Two-way ANOVA | | $F_{(1,18)}=1.253$ | | *p*=0.2777 | | |
| | Cell-type | | Two-way ANOVA | | $F_{(2,18)}=42.19$ | | *********p*<0.0001** | | |
| | **Post-test** | | | | | | | | |
| | P21 vs. P65 PV | | Šídák's multiple comparisons | | | | *p*=0.1897 | | |
| | P21 vs. P65 SST | | Šídák's multiple comparisons | | | | *p*=0.9998 | | |
| | P21 vs. P65 Other | | Šídák's multiple comparisons | | | | *p*=0.9998 | | |
| **K, L** | **Factor or comparison** | | **Statistical test** | | **F statistic** | | **P value** | | **Ns** |
| | P21 WT vs. KI | | Mann-Whitney test | | | | *p*=0.7857 | | WT *N*=5 at P21, KI N=3 at P21 |
| | P65 WT vs. KI | | Mann-Whitney test | | | | *p*=0.0952 | | WT N=5 at P65, KI N=5 at P65 |

**Figure 2. PV-specific mC4-OE causes an increase in anxiety-like behavior in male mice.**

| **Panel** | | | | | |
|---|---|---|---|---|---|
| **A** | N/A | | | | |
| **B** | N/A | | | | |
| **C** | Percent time spent in the open arms of the EZM did not differ between groups. | | | | |
| | **Factor or comparison** | **Statistical test** | **F statistic** | **P value** | **Ns** |
| | WT vs. KI | Mann-Whitney test | | *P*=0.1912 | |
| **D** | **Decreased time in the EZM open arms in male but not female KIs relative to WTs.** | | | | |
| | **Factor or comparison** | **Statistical test** | **F statistic** | **P value** | **Ns** |
| | Condition x Sex | Two-way ANOVA | F(1,34)=5.324 | ***p=0.0273** | |
| | Condition | Two-way ANOVA | F(1,34)=3.579 | *p*=0.0671 | |
| | Sex | Two-way ANOVA | F(1,34)=0.06062 | *p*=0.8070 | |
| | **Post-test** | | | | |
| | WT vs. KI males | Šídák's multiple comparisons | | ***p=0.0109** | |
| | WT vs. KI females | Šídák's multiple comparisons | | *p*=0.9474 | |
| **E** | **KI mice spent significantly less time in the light zone of the LDB compared to WT mice** | | | | |
| | **Factor or comparison** | **Statistical test** | **F statistic** | **P value** | **Ns** |
| | | t-test with Welch’s | | ***p=0.0130** | |
| **F** | **No sex-dependent differences in LDB light zone time between groups** | | | | |
| | **Factor or comparison** | **Statistical test** | **F statistic** | **P value** | **Ns** |
| | Condition x Sex | Two-way ANOVA | F(1,34)=0.3253 | *p*=0.5722 | |
| | Condition | Two-way ANOVA | F(1,34)=4.137 | ***p=0.0498** | |
| | Sex | Two-way ANOVA | F(1,34)=6.147 | ***p=0.0183** | |

| | | | | | |
|---|---|---|---|---|---|
| | **Post-test** | | | | |
| | WT vs. KI males | Šídák's multiple comparisons | | *p*=0.5211 | |
| | WT vs. KI females | Šídák's multiple comparisons | | *p*=0.1430 | |
| **G** | **Factor or comparison** | **Statistical test** | **F statistic** | **P value** | **Ns** |
| | Wt vs KI | unpaired t-test | unpaired t-test | | |
| **H** | **Significant increase in the latency to feed for KI males compared to WT controls.** | | | | |
| | **Factor or comparison** | **Statistical test** | **F statistic** | **P value** | **Ns** |
| | Condition x Sex | Two-way ANOVA | F(1,33)=6.718 | *p=0.0141 | |
| | Condition | Two-way ANOVA | $F_{(1,33)}=2.061$ | p=0.1606 | |
| | Sex | Two-way ANOVA | $F_{(1,33)}=0.4862$ | *p*=0.4905 | |
| | **Post-test** | | | | |
| | WT vs. KI males | Šídák's multiple comparisons | | *p=0.0159 | |
| | WT vs. KI females | Šídák's multiple comparisons | | *p*=0.6584 | |
| **I** | **Significant increase in the Z-Anxiety of KI mice relative to WTs.** | | | | |
| | **Factor or comparison** | **Statistical test** | **F statistic** | **P value** | **Ns** |
| | Wt vs KI | unpaired t-test | | *p=0.0114 | |
| **J** | **Compared to WT controls, there was a significant increase in the Z-Anxiety of KI male but not KI female mice** | | | | |
| | **Factor or comparison** | **Statistical test** | **F statistic** | **P value** | **Ns** |
| | Condition x Sex | Two-way ANOVA | $F_{(1,34)}=5.209$ | *p=0.0289 | |
| | Condition | Two-way ANOVA | F(1,34)=5.325 | *p=0.0272 | |
| | Sex | Two-way ANOVA | $F_{(1,34)}=5.756$ | *p=0.0221 | |
| | **Post-test** | | | | |
| | WT vs. KI males | Šídák's multiple comparisons | | **p=0.0053 | WT *N*=12 males, *N*=7 females |
| | WT vs. KI females | Šídák's multiple comparisons | | *p*=0.9998 | KI N=7 males, N=12 females |

| Figure 3. Increased levels of *mC4* in PV cells disrupts active but not passive social behaviors. | | | | | |
|---|---|---|---|---|---|
| **Panel** | | | | | |
| **A** | **N/A** | | | | |
| **B** | **KI mice explored the novel object as much as WT controls** | | | | |
| | **Factor or comparison** | **Statistical test** | **F statistic** | **P value** | **Ns** |
| | Condition x Sex | Two-way ANOVA | $F_{(1,44)}$=1.068 | *p*=0.3072 | |
| | Condition | Two-way ANOVA | $F_{(1,44)}$=0.008471 | *p*=0.9271 | |
| | Sex | Two-way ANOVA | $F_{(1,44)}$=0.02598 | *p*=0.8727 | |
| **C** | **N/A** | | | | |
| **D** | **Relative to WT controls, KI mice spent significantly less time engaged in the active snout-rear interaction type** | | | | |
| | **Factor or comparison** | **Statistical test** | **F statistic** | **P value** | **Ns** |
| | Wt vs KI | t-test with Welch's | | *******p*=0.0099** | **Ns** |
| | **There were no sex-related differences in active-snout-rear interaction between groups.** | | | | |
| | **Factor or comparison** | **Statistical test** | **F statistic** | **P value** | **Ns** |
| | Condition x Sex | Two-way ANOVA | $F_{(1,44)}$=0.007274 | *p*=0.9324 | |
| | Condition | Two-way ANOVA | $F_{(1,44)}$=7.127 | ******p*=0.0106** | |
| | Sex | Two-way ANOVA | **$F_{(1,44)}$=0.1121** | *p*=0.7394 | |
| | **Post-test** | | | | |
| | WT vs. KI males | Šídák's multiple comparisons | | *p*=0.1325 | |
| | WT vs. KI females | Šídák's multiple comparisons | | *p*=0.1218 | |
| **E** | **There was no significant change in Z-Sociability in either KI male or female mice** | | | | |
| | **Factor or comparison** | **Statistical test** | **F statistic** | **P value** | **Ns** |
| | Condition x Sex | Two-way ANOVA | $F_{(1,44)}$=1.808 | *p*=0.1857 | |
| | Condition | Two-way ANOVA | $F_{(1,44)}$=0.09738 | *p*=0.7565 | |
| | Sex | Two-way ANOVA | $F_{(1,44)}$=1.507 | *p*=0.2261 | |

| | | | | | | | | | |
|---|---|---|---|---|---|---|---|---|---|
| | | | | | | | | | WT, *N*=13 males, *N*=12 females |
| | | | | | | | | | KI, *N*=12 males, *N*=11 females |

| Figure 4. Sex-related difference in excitatory-inhibitory dynamics in mPFC PV cells with increased levels of *mC4* in PV cells. | | | | | |
|---|---|---|---|---|---|
| **Panel** | | | | | |
| **A** | **N/A** | | | | |
| **B** | **No significant change in mEPSC amplitude in KI mice, compared to controls** | | | | |
| | **Factor or comparison** | **Statistical test** | **F statistic** | **P value** | **Ns** |
| | Condition x Sex | Two-way ANOVA | $F_{(1,66)}$=1.111 | *p*=0.2956 | |
| | Condition | Two-way ANOVA | $F_{(1,66)}$=0.7845 | *p*=0.3790 | |
| | Sex | Two-way ANOVA | $F_{(1,66)}$=5.552 | ***p=0.0214** | |
| | **Post-test** | | | | |
| | WT vs. KI males | Šídák's multiple comparisons | | *p*=0.3225 | |
| | WT vs. KI females | Šídák's multiple comparisons | | *p*=0.9910 | |
| **C** | **No shift in the cumulative frequency distribution of mEPSC amplitudes in KI mice** | | | | |
| | **Factor or comparison** | **Statistical test** | **F statistic** | **P value** | **Ns** |
| | WT vs. KI males | Kolmogorov-Smirnov | | *p*=0.5758 | |
| | WT vs. KI females | Kolmogorov-Smirnov | | *p*=0.7810 | |
| **D** | ***mC4* OE led to a significant decrease in mEPSC frequency in KI male mice, but not KI female mice** | | | | |
| | **Factor or comparison** | **Statistical test** | **F statistic** | **P value** | **Ns** |
| | Condition x Sex | Two-way ANOVA | **$F_{(1,66)}$=2.138** | *p*=0.1485 | |
| | Condition | Two-way ANOVA | **$F_{(1,66)}$=8.979** | ****p=0.0038** | |
| | Sex | Two-way ANOVA | **$F_{(1,66)}$=0.3501** | *p*=0.5561 | |
| **E** | **Increased *mC4* expression caused a rightward shift in the distribution of mEPSC inter-event-intervals (IEIs) of PV cells in KI male mice, but not KI female mice** | | | | |
| | **Factor or comparison** | **Statistical test** | **F statistic** | **P value** | **Ns** |
| | WT vs. KI males | Kolmogorov-Smirnov | | ****p=0.0059** | |
| | WT vs. KI females | Kolmogorov-Smirnov | | *p*=0.3079 | |

| | | | | | | | | |
|---|---|---|---|---|---|---|---|---|
| **F** | **mEPSC Rise$_{10-90}$ was not impacted in KI mice, relative to WT controls** | | | | | | | |
| | **Factor or comparison** | | **Statistical test** | | **F statistic** | | **P value** | | **Ns** |
| | Condition x Sex | | Two-way ANOVA | | $F_{(1,66)}=0.8620$ | | *p*=0.3566 | | |
| | Condition | | Two-way ANOVA | | $F_{(1,66)}=0.02164$ | | *p*=0.8835 | | |
| | Sex | | Two-way ANOVA | | $F_{(1,66)}=1.121$ | | *p*=0.2935 | | |
| **G** | **mEPSC Decay$_{tau}$ was not changed in KI mice** | | | | | | | |
| | **Factor or comparison** | | **Statistical test** | | **F statistic** | | **P value** | | **Ns** |
| | Condition x Sex | | Two-way ANOVA | | $F_{(1,66)}=1.426$ | | *p*=0.2367 | | |
| | Condition | | Two-way ANOVA | | $F_{(1,66)}=0.9701$ | | *p*=0.3282 | | |
| | Sex | | Two-way ANOVA | | $F_{(1,66)}=2.700$ | | *p*=0.1051 | | |
| **H** | N/A | | | | | | | | |
| **I** | **mIPSC amplitude was significantly increased in KI male, but not KI female mice** | | | | | | | |
| | **Factor or comparison** | | **Statistical test** | | **F statistic** | | **P value** | | **Ns** |
| | Condition x Sex | | Two-way ANOVA | | $F_{(1,53)}=1.557$ | | *p*=0.2176 | | |
| | Condition | | Two-way ANOVA | | $F_{(1,53)}=5.831$ | | ***p*=0.0192** | | |
| | Sex | | Two-way ANOVA | | $F_{(1,53)}=0.2637$ | | *p*=0.6097 | | |
| | **Post-test** | | | | | | | | |
| | WT vs. KI males | | Šídák's multiple comparisons | | | | ***p*=0.0232** | | |
| | WT vs. KI females | | Šídák's multiple comparisons | | | | *p*=0.6599 | | |
| **J** | **Increased expression of *mC4* caused a rightward shift in the distribution of mIPSC amplitudes in KI male mice** | | | | | | | |
| | **Factor or comparison** | | **Statistical test** | | **F statistic** | | **P value** | | **Ns** |
| | WT vs. KI males | | Kolmogorov-Smirnov | | | | ***p*=0.0404** | | |
| | WT vs. KI females | | Kolmogorov-Smirnov | | | | *p*=0.9048 | | |
| **K** | **mIPSC frequency was not changed in KI mice** | | | | | | | |
| | **Factor or comparison** | | **Statistical test** | | **F statistic** | | **P value** | | **Ns** |
| | Condition x Sex | | Two-way ANOVA | | $F_{(1,53)}=1.117$ | | *p*=0.2954 | | |

| | | | | | | | | | |
|---|---|---|---|---|---|---|---|---|---|
| | Condition | | Two-way ANOVA | | $F_{(1,53)}$=0.04095 | | *p*=0.8404 | | |
| | Sex | | Two-way ANOVA | | $F_{(1,53)}$=0.5669 | | *p*=0.4548 | | |
| **L** | **No shift in the distribution of mIPSC IEIs of KI mice** | | | | | | | | |
| | **Factor or comparison** | | **Statistical test** | | **F statistic** | | **P value** | | **Ns** |
| | WT vs. KI males | | Kolmogorov-Smirnov | | | | *p*=0.4149 | | |
| | WT vs. KI females | | Kolmogorov-Smirnov | | | | *p*=0.9988 | | |
| **M** | **Relative to controls, there were no changes in mIPSC Rise$_{10\text{-}90}$ with increased levels of *mC4* in PV cells** | | | | | | | | |
| | **Factor or comparison** | | **Statistical test** | | **F statistic** | | **P value** | | **Ns** |
| | Condition x Sex | | Two-way ANOVA | | $F_{(1,53)}$=7.666 | | ****p=0.0077** | | |
| | Condition | | Two-way ANOVA | | $F_{(1,53)}$=0.2031 | | *p*=0.6541 | | |
| | Sex | | Two-way ANOVA | | $F_{(1,53)}$=0.4950 | | *p*=0.4848 | | |
| | **Post-test** | | | | | | | | |
| | WT vs. KI males | | Šídák's multiple comparisons | | | | *p*=0.0507 | | |
| | WT vs. KI females | | Šídák's multiple comparisons | | | | *p*=0.2079 | | |
| **N** | **mIPSC Decay$_{tau}$ was not changed in KI mice** | | | | | | | | |
| | Condition x Sex | | Two-way ANOVA | | $F_{(1,53)}$=1.708 | | *p*=0.1969 | | |
| | Condition | | Two-way ANOVA | | $F_{(1,53)}$=0.01747 | | *p*=0.8953 | | |
| | Sex | | Two-way ANOVA | | $F_{(1,53)}$=0.1166 | | *p*=0.7341 | | |
| | | | | | | | | | mEPSC WT, *N*=20 males, *N*=17 females |
| | | | | | | | | | mEPSC KI: *N*=15 males, *N*=18 females |
| | | | | | | | | | mIPSC WT *N*=14 males, *N*=14 females |
| | | | | | | | | | mIPSC KI: *N*=15 males, *N*=14 females |

| | | | | | |
|---|---|---|---|---|---|
| **Figure 5. PV-specific mC4-OE leads to opposing changes in excitability of PV cells in male and female mice.** | | | | | |
| **Panel** | | | | | |
| **A** | N/A | | | | |
| **B** | **PV cells in KI male mice spike less than PV cells in WT male mice** | | | | |
| | **Factor or comparison** | **Statistical test** | **F statistic** | **P value** | **Ns** |
| | Condition x Current | RM Two-way ANOVA | $F_{(24,840)}=1.941$ | *******p*=0.0045** | |
| | Condition | RM Two-way ANOVA | $F_{(1,35)}=1.989$ | *p*=0.1672 | |
| | Current | RM Two-way ANOVA | $F_{(1.656,57.95)}=416.4$ | *********p*<0.0001** | |
| **C** | ***mC4* OE led to an increase in the excitability of PV cells in KI female mice, relative to controls** | | | | |
| | **Factor or comparison** | **Statistical test** | **F statistic** | **P value** | **Ns** |
| | Condition x Current | RM Two-way ANOVA | $F_{(24,816)}=3.428$ | *********p*<0.0001** | |
| | Condition | RM Two-way ANOVA | $F_{(1,34)}=4.178$ | ******p*=0.0488** | |
| | Current | RM Two-way ANOVA | $F_{(1.719,58.46)}=568.3$ | *********p*<0.0001** | |
| **D** | **Rheobase was decreased in KI female mice, relative to controls** | | | | |
| | **Factor or comparison** | **Statistical test** | **F statistic** | **P value** | **Ns** |
| | Condition x Sex | Two-way ANOVA | $F_{(1,69)}=10.23$ | *******p*=0.0021** | |
| | Condition | Two-way ANOVA | $F_{(1,69)}=0.008666$ | *p*=0.9261 | |
| | Sex | Two-way ANOVA | $F_{(1,69)}=2.932$ | *p*=0.0913 | |
| | **Post-test** | | | | |
| | WT vs. KI males | Šídák's multiple comparisons | | *p*=0.0596 | |
| | WT vs. KI females | Šídák's multiple comparisons | | ******p*=0.0472** | |
| **E** | **PV-mC4-OE drove a shift in PV cell resting membrane voltage towards a more depolarized $V_m$** | | | | |
| | **Factor or comparison** | **Statistical test** | **F statistic** | **P value** | **Ns** |
| | WT vs. KI | t-test with Welch’s | | ******p*=0.0356** | |
| **F** | N/A | | | | |

| | | | | | |
|---|---|---|---|---|---|
| **G** | **OE of *mC4* decreased the spike frequency of PYRs in KI male mice, relative to controls** | | | | |
| | **Factor or comparison** | **Statistical test** | **F statistic** | **P value** | **Ns** |
| | Condition x Current | RM Two-way ANOVA | $F_{(24,696)}$=1.975 | ****p=0.0038** | |
| | Condition | RM Two-way ANOVA | $F_{(1,29)}$=2.646 | *p*=0.1147 | |
| | Current | RM Two-way ANOVA | $F_{(1.350,39.15)}$=619.8 | ******p<0.0001** | |
| **H** | **No differences in the excitability of PYRs in females between groups** | | | | |
| | Condition x Current | RM Two-way ANOVA | $F_{(24,816)}$=0.2813 | *p*=0.9998 | |
| | Condition | RM Two-way ANOVA | $F_{(1,34)}$=0.1154 | *p*=0.7361 | |
| | Current | RM Two-way ANOVA | $F_{(1.467,49.89)}$=810.0 | ******p<0.0001** | |
| **I** | **No changes in PYR Rheobase** | | | | |
| | **Factor or comparison** | **Statistical test** | **F statistic** | **P value** | **Ns** |
| | Condition x Sex | Two-way ANOVA | $F_{(1,63)}$=5.148 | ***p=0.0267** | |
| | Condition | Two-way ANOVA | $F_{(1,63)}$=0.2332 | *p*=0.6308 | |
| | Sex | Two-way ANOVA | $F_{(1,63)}$=0.8369 | *p*=0.3638 | |
| | **Post-test** | | | | |
| | WT vs. KI males | Šídák's multiple comparisons | | *p*=0.1262 | |
| | WT vs. KI females | Šídák's multiple comparisons | | *p*=0.3502 | |
| **J** | **No change overall in PYR resting $V_m$ in KI mice, compared to controls** | | | | |
| | **Factor or comparison** | **Statistical test** | **F statistic** | **P value** | **Ns** |
| | WT vs. KI | Mann-Whitney test | | *p*=0.6464 | |
| | | | | | PV cell WT *N*=19 males, *N*=17 females |
| | | | | | PV cell KI: *N*=18 males, *N*=19 females |
| | | | | | PYR WT *N*=15 males, *N*=18 females |
| | | | | | PYR KI: *N*=16 males, *N*=18 females |

| **Figure 6. No changes in anxiety-like behavior with pan-neuronal overexpression of *mC4*.** | | | | | | | | | |
|---|---|---|---|---|---|---|---|---|---|
| **Panel** | | | | | | | | | |
| **A** | N/A | | | | | | | | |
| **B** | **Pan-neuronal *mC4* OE does not alter locomotion** | | | | | | | | |
| | **Factor or comparison** | | **Statistical test** | | **F statistic** | | **P value** | | **Ns** |
| | WT vs. KI | | t-test | | | | *p*=0.9092 | | |
| **C** | **Pan-neuronal *mC4* OE does not alter locomotion** | | | | | | | | |
| | **Factor or comparison** | | **Statistical test** | | **F statistic** | | **P value** | | **Ns** |
| | Condition x Sex | | Two-way ANOVA | | $F_{(1,36)}$=1.039 | | *p*=0.3148 | | |
| | Condition | | Two-way ANOVA | | $F_{(1,36)}$=0.00415 | | *p*=0.9490 | | |
| | Sex | | Two-way ANOVA | | $F_{(1,36)}$=0.4105 | | *p*=0.5258 | | |
| **D** | **No change in time spent in the open arms with pan-neuronal *mC4* OE** | | | | | | | | |
| | **Factor or comparison** | | **Statistical test** | | **F statistic** | | **P value** | | **Ns** |
| | WT vs. KI | | Mann-Whitney test | | | | *p*=0.6487 | | |
| **E** | **No change in time spent in the open arms with pan-neuronal *mC4* OE** | | | | | | | | |
| | **Factor or comparison** | | **Statistical test** | | **F statistic** | | **P value** | | **Ns** |
| | Condition x Sex | | Two-way ANOVA | | $F_{(1,36)}$=1.225 | | *p*=0.2757 | | |
| | Condition | | Two-way ANOVA | | $F_{(1,36)}$=0.1021 | | *p*=0.7512 | | |
| | Sex | | Two-way ANOVA | | $F_{(1,36)}$=0.00008 | | *p*=0.9928 | | |
| **F** | **Time spent in the light zone was not altered by pan-neuronal *mC4* OE** | | | | | | | | |
| | **Factor or comparison** | | **Statistical test** | | **F statistic** | | **P value** | | **Ns** |
| | WT vs. KI | | Mann-Whitney test | | | | *p*=0.9360 | | |
| **G** | **Time spent in the light zone was not altered by pan-neuronal *mC4* OE** | | | | | | | | |

| | Factor or comparison | | Statistical test | | F statistic | | P value | | Ns |
|---|---|---|---|---|---|---|---|---|---|
| | Condition x Sex | | Two-way ANOVA | | $F_{(1,36)}$=4.620 | | ***p*=0.0384** | | |
| | Condition | | Two-way ANOVA | | $F_{(1,36)}$=0.07082 | | *p*=0.7917 | | |
| | Sex | | Two-way ANOVA | | $F_{(1,36)}$=2.552 | | *p*=0.1189 | | |
| | **Post-test** | | | | | | | | |
| | WT vs. KI males | | Šídák's multiple comparisons | | | | *p*=0.1977 | | |
| | WT vs. KI females | | Šídák's multiple comparisons | | | | *p*=0.3281 | | |
| **H** | **OE of *mC4* in neurons did not alter feed latency** | | | | | | | | |
| | **Factor or comparison** | | **Statistical test** | | **F statistic** | | **P value** | | **Ns** |
| | WT vs. KI | | t-test | | | | *p*=0.1761 | | |
| **I** | **OE of *mC4* in neurons did not alter feed latency** | | | | | | | | |
| | **Factor or comparison** | | **Statistical test** | | **F statistic** | | **P value** | | **Ns** |
| | Condition x Sex | | Two-way ANOVA | | $F_{(1,29)}$=4.093 | | *p*=0.0524 | | |
| | Condition | | Two-way ANOVA | | $F_{(1,29)}$=1.328 | | *p*=0.2586 | | |
| | Sex | | Two-way ANOVA | | $F_{(1,29)}$=0.4055 | | *p*=0.5293 | | |
| **J** | ***mC4* OE in neurons did not lead to changes in overall anxiety-like behavior.** | | | | | | | | |
| | **Factor or comparison** | | **Statistical test** | | **F statistic** | | **P value** | | **Ns** |
| | WT vs. KI | | t-test with Welch's | | | | *p*=0.5358 | | |
| **K** | ***mC4* OE in neurons did not lead to changes in overall anxiety-like behavior.** | | | | | | | | |
| | **Factor or comparison** | | **Statistical test** | | **F statistic** | | **P value** | | **Ns** |
| | Condition x Sex | | Two-way ANOVA | | $F_{(1,36)}$=0.3797 | | *p*=0.5416 | | |
| | Condition | | Two-way ANOVA | | $F_{(1,36)}$=0.03077 | | *p*=0.8617 | | |
| | Sex | | Two-way ANOVA | | $F_{(1,36)}$=1.008 | | *p*=0.3221 | | |
| | | | | | | | | | WT *N*=9 males, *N*=10 females |
| | | | | | | | | | KI: *N*=10 males, *N*=11 females |

| | | | | | |
|---|---|---|---|---|---|
| **Figure 7. Disrupted neural communication and hyperexcitability in a network model of male mice with increased levels of *mC4* in PV cells.** | | | | | |
| **Panel** | | | | | |
| **A** | N/A | | | | |
| **B** | **The firing rate (FR) of PV2 in WT and KI networks at peak *Iapp* of 200 pA** | | | | |
| | **Factor or comparison** | **Statistical test** | **F statistic** | **P value** | **Ns** |
| | Condition x Sex, 200 pA | Two-way ANOVA | $F_{(1,56)}$=49.79 | ****$\boldsymbol{p<0.0001}$ | |
| | Condition, 200 pA | Two-way ANOVA | $F_{(1,56)}$=54.23 | ****$\boldsymbol{p<0.0001}$ | |
| | Sex, 200 pA | Two-way ANOVA | $F_{(1,56)}$=0.7170 | $p$=0.4007 | |
| | **Post-test** | | | | |
| | WT vs. KI males, 200 pA | Šídák's multiple comparisons | | ****$\boldsymbol{p<0.0001}$ | |
| | WT vs. KI females, 200 pA | Šídák's multiple comparisons | | $p$=0.9706 | |
| | **The firing rate (FR) of PV2 in WT and KI networks at peak *Iapp* of 275 pA** | | | | |
| | **Factor or comparison** | **Statistical test** | **F statistic** | **P value** | **Ns** |
| | Condition x Sex, 275 pA | Two-way ANOVA | $F_{(1,56)}$=71.87 | ****$\boldsymbol{p<0.0001}$ | |
| | Condition, 275 pA | Two-way ANOVA | $F_{(1,56)}$=58.30 | ****$\boldsymbol{p<0.0001}$ | |
| | Sex, 275 pA | Two-way ANOVA | $F_{(1,56)}$=1.390 | $p$=0.2434 | |
| | **Post-test** | | | | |
| | WT vs. KI males, 275 pA | Šídák's multiple comparisons | | ****$\boldsymbol{p<0.0001}$ | |
| | WT vs. KI females, 275 pA | Šídák's multiple comparisons | | $p$=0.8010 | |
| | **The firing rate (FR) of PV2 in WT and KI networks at peak *Iapp* of 350 pA** | | | | |
| | **Factor or comparison** | **Statistical test** | **F statistic** | **P value** | **Ns** |
| | Condition x Sex, 350 pA | Two-way ANOVA | $F_{(1,56)}$=114.7 | ****$\boldsymbol{p<0.0001}$ | |
| | Condition, 350 pA | Two-way ANOVA | $F_{(1,56)}$=47.97 | ****$\boldsymbol{p<0.0001}$ | |

| | | | | | | | | | |
|---|---|---|---|---|---|---|---|---|---|
| | Sex, 350 pA | | Two-way ANOVA | | $F_{(1,56)}$=3.473 | | *p*=.0676 | | |
| | **Post-test** | | | | | | | | |
| | WT vs. KI males, 350 pA | | Šídák's multiple comparisons | | | | *********p*<0.0001** | | |
| | WT vs. KI females, 350 pA | | Šídák's multiple comparisons | | | | ******p*=0.0195** | | |
| **C** | **The FR of PYR2 in WT and KI networks at peak *Iapp* of 200 pA** | | | | | | | | |
| | **Factor or comparison** | | **Statistical test** | | **F statistic** | | **P value** | | **Ns** |
| | Condition x Sex, 200 pA | | Two-way ANOVA | | $F_{(1,56)}$=98.21 | | *********p*<0.0001** | | |
| | Condition, 200 pA | | Two-way ANOVA | | $F_{(1,56)}$=24.10 | | *********p*<0.0001** | | |
| | Sex, 200 pA | | Two-way ANOVA | | $F_{(1,56)}$=113.4 | | *********p*<0.0001** | | |
| | **Post-test** | | | | | | | | |
| | WT vs. KI males, 200 pA | | Šídák's multiple comparisons | | | | *********p*<0.0001** | | |
| | WT vs. KI females, 200 pA | | Šídák's multiple comparisons | | | | *******p*=0.0016** | | |
| | **The FR of PYR2 in WT and KI networks at peak *Iapp* of 275 pA** | | | | | | | | |
| | **Factor or comparison** | | **Statistical test** | | **F statistic** | | **P value** | | **Ns** |
| | Condition x Sex, 275 pA | | Two-way ANOVA | | $F_{(1,56)}$=74.29 | | *********p*<0.0001** | | |
| | Condition, 275 pA | | Two-way ANOVA | | $F_{(1,56)}$=2.972 | | *p*=0.0902 | | |
| | Sex, 275 pA | | Two-way ANOVA | | $F_{(1,56)}$=163.2 | | *********p*<0.0001** | | |
| | **Post-test** | | | | | | | | |
| | WT vs. KI males, 275 pA | | Šídák's multiple comparisons | | | | *********p*<0.0001** | | |
| | WT vs. KI females, 275 pA | | Šídák's multiple comparisons | | | | *********p*<0.0001** | | |
| | **The FR of PYR2 in WT and KI networks at peak *Iapp* of 350 pA** | | | | | | | | |
| | **Factor or comparison** | | **Statistical test** | | **F statistic** | | **P value** | | **Ns** |
| | Condition x Sex, 350 pA | | Two-way ANOVA | | $F_{(1,56)}$=44.10 | | *********p*<0.0001** | | |

| | | | | | | | | | |
|---|---|---|---|---|---|---|---|---|---|
| | Condition, 350 pA | | Two-way ANOVA | | $F_{(1,56)}$=0.1087 | | *p*=0.7429 | | |
| | Sex, 350 pA | | Two-way ANOVA | | $F_{(1,56)}$=162.9 | | ****p<0.0001** | | |
| | **Post-test** | | | | | | | | |
| | WT vs. KI males, 350 pA | | Šídák's multiple comparisons | | | | ****p<0.0001** | | |
| | WT vs. KI females, 350 pA | | Šídák's multiple comparisons | | | | ****p<0.0001** | | |
| **D** | **Transfer entropy (TE) at each delay (lag) of PYR1 onto PYR2 in male WT (*blue*) and male KI (*red*) networks at peak *Iapp* of 200 pA** | | | | | | | | |
| | **Factor or comparison** | | **Statistical test** | | **F statistic** | | **P value** | | **Ns** |
| | Condition x Lag, 200 pA | | Two-way ANOVA | | $F_{(7,196)}$=15.72 | | ****p<0.0001** | | |
| | Condition, 200 pA | | Two-way ANOVA | | $F_{(1,28)}$=45.51 | | ****p<0.0001** | | |
| | Lag, 200 pA | | Two-way ANOVA | | $F_{(2.031,56.86)}$=172.5 | | ****p<0.0001** | | |
| | **Transfer entropy (TE) at each delay (lag) of PYR1 onto PYR2 in male WT (*blue*) and male KI (*red*) networks at peak *Iapp* of 275 pA** | | | | | | | | |
| | **Factor or comparison** | | **Statistical test** | | **F statistic** | | **P value** | | **Ns** |
| | Condition x Lag, 275 pA | | Two-way ANOVA | | $F_{(7,196)}$=8.938 | | ****p<0.0001** | | |
| | Condition, 275 pA | | Two-way ANOVA | | $F_{(1,28)}$=38.25 | | ****p<0.0001** | | |
| | Lag, 275 pA | | Two-way ANOVA | | $F_{(2.768, 77.52)}$=223.9 | | ****p<0.0001** | | |
| | **Transfer entropy (TE) at each delay (lag) of PYR1 onto PYR2 in male WT (*blue*) and male KI (*red*) networks at peak *Iapp* of 350pA** | | | | | | | | |
| | **Factor or comparison** | | **Statistical test** | | **F statistic** | | **P value** | | **Ns** |
| | Condition x Lag, 350 pA | | Two-way ANOVA | | $F_{(7,196)}$=9.861 | | ****p<0.0001** | | |
| | Condition, 350 pA | | Two-way ANOVA | | $F_{(1,28)}$=32.48 | | ****p<0.0001** | | |
| | Lag, 350 pA | | Two-way ANOVA | | $F_{(2.687,75.22)}$=149.7 | | ****p<0.0001** | | |
| **E** | **TE at each delay (lag) of PYR1 onto PYR2 in female WT (*blue*) and female KI (*red*) networks at peak *Iapp* of 200 pA** | | | | | | | | |
| | **Factor or comparison** | | **Statistical test** | | **F statistic** | | **P value** | | **Ns** |
| | Condition x Lag, 200 pA | | Two-way ANOVA | | $F_{(7,196)}$=1.576 | | *p*=0.1446 | | |
| | Condition, 200 pA | | Two-way ANOVA | | $F_{(1,28)}$=6.672 | | ***p=0.0153** | | |
| | Lag, 200 pA | | Two-way ANOVA | | $F_{(1.772,49.61)}$=121.6 | | ****p<0.0001** | | |

| | | | | | | | | | |
|---|---|---|---|---|---|---|---|---|---|
| | **TE at each delay (lag) of PYR1 onto PYR2 in female WT (*blue*) and female KI (*red*) networks at peak *Iapp* of 275 pA** | | | | | | | | |
| | **Factor or comparison** | | **Statistical test** | | **F statistic** | | **P value** | | **Ns** |
| | Condition x Lag, 275 pA | | Two-way ANOVA | | $F_{(7,196)}=0.4033$ | | *p*=0.8995 | | |
| | Condition, 275 pA | | Two-way ANOVA | | $F_{(1,28)}=14.14$ | | ********p*=0.0008** | | |
| | Lag, 275 pA | | Two-way ANOVA | | $F_{(1.484,41.55)}=127.9$ | | *********p*<0.0001** | | |
| | **TE at each delay (lag) of PYR1 onto PYR2 in female WT (*blue*) and female KI (*red*) networks at peak *Iapp* of 350 pA** | | | | | | | | |
| | **Factor or comparison** | | **Statistical test** | | **F statistic** | | **P value** | | **Ns** |
| | Condition x Lag, 350 pA | | Two-way ANOVA | | $F_{(7,196)}=3.124$ | | *******p*=0.0038** | | |
| | Condition, 350 pA | | Two-way ANOVA | | $F_{(1,28)}=13.92$ | | ********p*=0.0009** | | |
| | Lag, 350 pA | | Two-way ANOVA | | $F_{(1.985,55.57)}=135.6$ | | *********p*<0.0001** | | |
| | | | | | | | | | WT network, *N*=15 |
| | | | | | | | | | KI network, *N*=15 |
